# Pansoma, a machine learning tool for identifying somatic variants using pangenome graphs

**DOI:** 10.64898/2026.05.27.726245

**Authors:** Jiawei Shen, Qichen Fu, Juan F. Macias-Velasco, Daofeng Li, Ting Wang

## Abstract

Somatic variant calling, the identification of mutations in non-germline cells acquired over an individual’s lifetime, is critical for studying diseases, including cancer, and for developing precision oncology strategies. Traditional somatic variant calling methods rely on linear reference genomes, which do not adequately capture human genetic diversity and result in reference bias, compromising the accuracy of somatic variant detection. Recently developed graph-based human pangenome reference represents diverse genetic variants across human populations and has promised to drive advances in many genetics and genomics studies. In this study, we introduced Pansoma, a novel pangenome-native and machine learning-based tool specifically designed for somatic variant calling using a pangenome graph reference. Pansoma performs somatic variant detection from both short- and long-read sequencing data by learning tensor representations of alignment on graph nodes rather than on a linear reference. Pansoma outputs variant representations anchored to the pangenome graph paths and conventional somatic variant calls remapped to the linear reference. Additionally, we provide accompanying bioinformatics tools tailored for graph-based genomic data management and variant calling results analysis. Benchmarking shows that Pansoma not only improves tumor-only somatic variant detection but also preserves graph-specific variant representations that are not directly recoverable from linear-reference outputs.

## Introduction

The identification of genetic variants from next-generation sequencing (NGS) data, a process known as variant calling, is a fundamental task in modern genomics, enabling breakthroughs in fields ranging from population genetics to the study of inherited diseases [1, 2]. A particularly critical application of this technology is the detection of somatic mutations, genetic alterations that are acquired during an organism’s lifetime rather than inherited. These mutations are the core drivers of tumor initiation and evolution [3–5]. Therefore, the accurate identification of somatic variants from NGS data is crucial for understanding tumor biology, enabling clinical diagnostics, and guiding targeted therapies [6, 7]. Over the past decade, numerous computational somatic variant-calling methods have been developed to address this challenge, primarily in the context of short-read sequencing data. These methods can generally be categorized into two main paradigms: tumor-normal paired analysis and tumor-only analysis. With the growing adoption of long-read sequencing technologies, somatic variant calling is now extending to data types that better resolve repetitive and complex genomic regions but also introduce distinct error profiles and alignment challenges. Tumor-normal paired analysis is considered the “gold standard” for somatic variant calling. By comparing sequencing data from a tumor and a matched normal tissue from the same individual, researchers can effectively filter out germline variants, thereby obtaining a high confidence set of somatic mutations. A suite of sophisticated tools, including GATK Mutect2 [8], VarScan2 [9], Strelka2 [10], and the deep-learning-based DeepSomatic [11], represent development for this tumor-normal paradigm.

However, in many clinical settings, acquiring high-quality matched normal samples is not feasible due to sample contamination, cost constraints, or ethical issues. Consequently, a vast amount of clinical tumor sequencing data consists only of tumor samples, creating an urgent need for high-precision tumor-only analysis methods. Detecting somatic mutations from tumor-only data is a formidable bioinformatics challenge. The core difficulty lies in distinguishing true somatic mutations from the vast background of germline variants without a matched normal sample for reference. There is a major challenge: it fails to filter out rare or private germline variants not cataloged in curated databases [12–15] and cannot reliably distinguish alignment artifacts from true somatic variants, leading to false positives. While many of these somatic variant calling tools can be adapted for a tumor-only workflow, their performance is often suboptimal due to the lack of a matched normal sample for direct germline filtering. Traditional methods such as Mutect2 [8] and Strelka [10] use heuristic filters and population databases, whereas recent AI-based methods like DeepSomatic [11] and ClairS [16] leverage deep learning to better distinguish somatic from germline variation, though both methods remain limited by linear reference genomes. These methods have improved the accuracy of tumor-only analysis to some extent, but there remains significant room for improvement.

Recently, the advent of the human pangenome reference has offered a transformative new resource for genomics [17–19]. Unlike a single linear reference genome, a pangenome is constructed from the genomes of multiple individuals from diverse backgrounds and represents a much broader spectrum of human genetic variation as a genome graph. This graph-based structure contains a multitude of alternative paths and a comprehensive catalog of known germline variants. Critically, modern alignment tools [20, 21] can now map raw sequencing data directly onto these complex pangenome graphs. The resulting alignment files explicitly encode the specific path or haplotype each read aligns to. For example, a read supporting a known germline variant will align preferentially, with a higher score, to the specific graph path containing that variant. This provides alignment-level evidence that the read sequence is consistent with a variant path already represented in the pangenome, which can help reduce misclassification of common germline variation as somatic mutations in tumor-only analysis. This proactive, evidence-based identification of germline variants during the alignment stage could offer a more robust foundation for tumor-only based somatic variant calling.

Here, we present Pansoma, a novel pangenome-native somatic variant caller specifically designed for tumor-only analysis directly on the pangenome graph. Our approach employs a pipeline that uses a deep convolutional neural network (CNN) to process graph alignments without projecting them onto a linear reference. To support this, we introduce a new node pileup (NPU) format, which stores graph-based alignments indexed by node ID for fast queries. Pansoma then constructs image-like tensors directly from the graph nodes, incorporating features such as base pairs, read quality, and CIGAR operations. Pansoma leverages these tensors to learn the characteristic patterns of somatic mutations and accurately predict candidate variants. This design allows it to fully exploit the richness of pangenome data while distinguishing somatic from germline variation, because germline variants represented in the pangenome can align cleanly to existing graph paths and are therefore less likely to be misclassified as novel variants. As a graph-native caller, Pansoma reports somatic variants using node IDs as positional references rather than linear coordinates. However, since the ecosystem of pangenome alignment processing is still maturing, we provide a projection to transform these graph-based alignments into a linear reference coordinate system for linear-reference benchmarking pipelines. Additionally, to retain variants that cannot be projected to the linear reference, we preserve graph-based variants using node positions to provide complementary information. We benchmarked Pansoma using datasets from two well-established community resources: the SMaHT consortium [22–24] and the Genome in a Bottle (GIAB) consortium [25]. The SMaHT consortium provides deeply characterized cancer-related sequencing resources, including the COLO829 melanoma cell line, which is widely used as a reference dataset for somatic variant benchmarking. GIAB provides high-confidence genomic benchmark datasets, including HG008 [25], which has recently been extended to support somatic variant evaluation. Using the COLO829 and HG008 datasets, we demonstrate that Pansoma achieves leading precision with comparable F1 scores in tumor-only somatic variant calling.

## Results

### Overview of Pansoma

Pansoma is a pangenome-native machine learning framework for tumor-only somatic variant calling from both short- and long-read sequencing data. Unlike conventional workflows that align reads to a single linear reference and infer variants only in linear coordinates, Pansoma performs variant discovery directly from graph alignment. This design allows read features to be interpreted on both the GRCh38 reference path and alternative haplotype paths in the pangenome graph, providing population-aware context for distinguishing somatic mutations from germline variation and alignment artifacts.

The Pansoma workflow consists of three major stages: graph-based data preparation, machine learning–based variant inference, and variant reconstruction with VCF generation. First (Fig.1a), sequencing reads from benchmark tumor datasets SMaHT COLO829 and HapMap [22, 24] and HG008 [26, 27] are aligned to the human pangenome graph using a graph-aware mapper, producing graph alignment files that retain the graph paths traversed by each read. These graph alignments are converted into node-level pileups, in which read segments traversing the same graph node are grouped together to summarize local read support at that node. Candidate variant sites are then identified from these graph-node pileups and transformed into image-like tensors (Fig. 1b). In each tensor, the horizontal axis represents the local sequence context around the candidate site, the vertical axis represents the stack of reads supporting that region, and separate channels encode read bases, base quality, mapping quality, mismatch flag, and CIGAR operation features (Methods). This representation enables the model to learn somatic-variant signals directly from graph-aligned read features for downstream training and inference.

**Figure 1.**
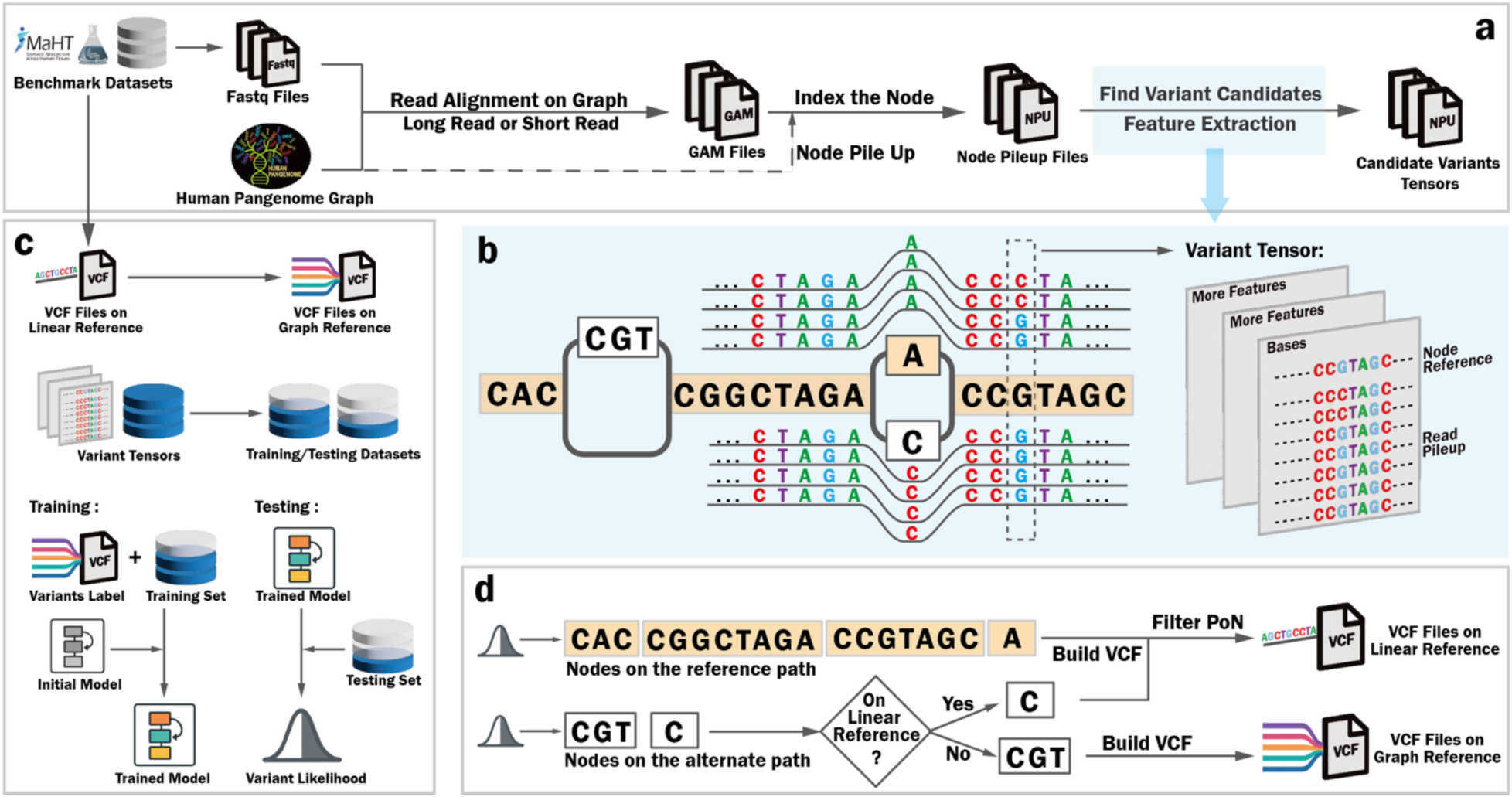
Overview of the Pansoma framework. (a) Sequencing reads from benchmark datasets are aligned to the human pangenome graph to generate graph alignment (GAM) files. Node-level pileups are then indexed to produce node pileup (NPU) files, from which candidate variant tensors are extracted for downstream feature analysis. (b) Schematic of graph-based tensor construction. Read pileups spanning reference and alternate graph nodes are converted into image-like variant tensors, with channels encoding base composition and additional features derived from the node reference and aligned reads. (c) Training and testing workflow. Benchmark VCFs defined on the linear reference are projected onto the graph coordinates, and together with tensorized candidate variants are used to assemble labeled training and testing datasets. These datasets are used to train the Pansoma model and evaluate variant likelihood predictions. (d) Post-processing and VCF generation. Predicted variants are reconstructed from graph nodes; variants on the linear reference path are reported as linear-reference VCF records, whereas variants on alternate graph paths are retained as graph-reference VCF records, followed by panel-of-normals filtering for linear-reference calls.

In the machine learning stage (Fig. 1c), Pansoma uses these graph-derived tensors as input to a convolutional neural network trained to classify candidate sites as somatic or non-somatic. Because available benchmark truth sets are primarily defined on the linear reference, benchmark VCF records are projected onto the pangenome graph to generate graph-coordinate labels for supervised training and evaluation. This strategy enables supervised learning in graph coordinates while leveraging established linear-reference truth sets.

Using the trained model, Pansoma performs somatic variant inference, after which the predicted variants are reconstructed from graph coordinates and reported in two complementary formats: linear-reference variants and graph-reference variants (Fig. 1d and Methods). Variants that can be projected to the GRCh38 reference path are converted into conventional linear-reference VCF records, allowing direct comparison with existing somatic variant callers and benchmark truth sets. Graph variants that cannot be unambiguously projected to GRCh38 are retained as graph- coordinate VCF records. This dual-output design preserves compatibility with existing VCF-based workflows while retaining somatic variants that may be lost in a purely linear-reference representation. Finally, because current PoN resources are defined in linear-reference coordinates, linear-reference calls are filtered using panel-of-normals resources to reduce residual germline and recurrent background variants.

### Graph-native candidate representation for Pansoma

A major challenge in developing Pansoma was constructing training and inference data directly from a pangenome graph. Although many somatic variant benchmark datasets are available in linear-reference coordinates, they do not directly provide graph-native representations suitable for machine learning. To address this, we developed a graph-based tensor construction strategy that uses graph nodes as the basic unit of local feature aggregation. After reads are aligned to the pangenome graph, Pansoma scans graph nodes and prioritizes regions supported by reads containing mismatches, indels or other non-perfect alignment patterns relative to the traversed graph node (Fig. 2a). These nodes are treated as potential variant-containing regions, and a local scan window is applied across each candidate node. Conceptually, this resembles a short linear pileup window, but the coordinate system is defined by graph nodes rather than by a single reference chromosome.

**Figure 2.**
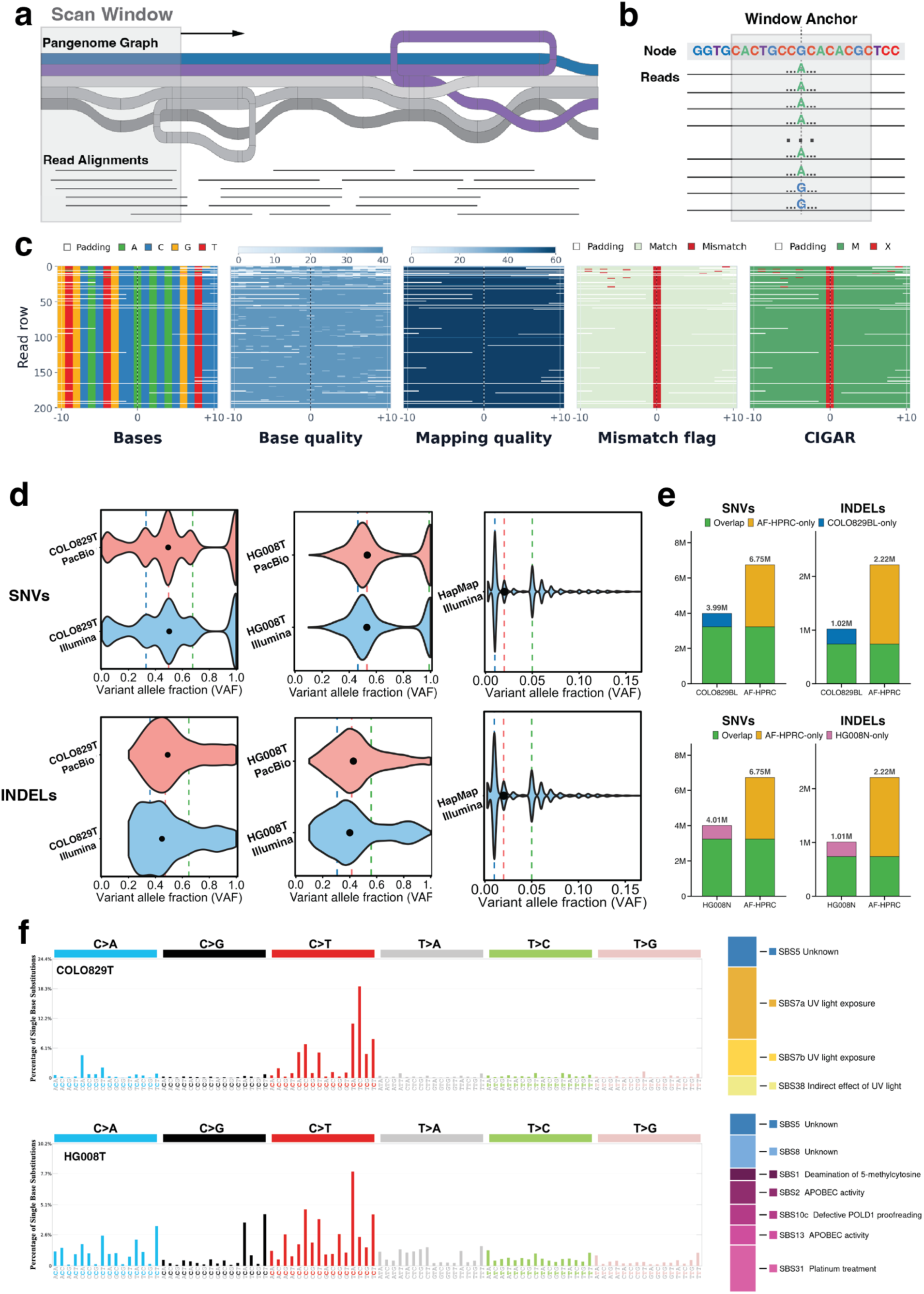
Benchmark resources and graph-native tensorization strategy for tumor-only somatic variant calling. (a) Schematic overview of graph-window scanning. Pansoma scans the pangenome graph and collects read alignments overlapping candidate regions. (b) Window anchoring strategy for candidate representation. A local window is centered around the candidate site within a graph node, and overlapping reads are aligned relative to the anchor. (c) Multi-channel pileup tensor used for model prediction. Each candidate is represented using five channels: read base, base quality, mapping quality, mismatch flag, and CIGAR operation across reads and local graph positions. (d) Variant allele fraction distributions of benchmark somatic SNVs and INDELs in COLO829T, HG008T, and HapMap mixture data. COLO829T and HG008T are shown for Illumina and PacBio platforms, whereas HapMap illustrates a low-VAF synthetic mixture setting. Black dots indicate median VAFs, and dashed vertical lines mark global VAF quartiles (25%, 50%, 75%). (e) Overlap between germline variants from the COLO829BL and HG008N normal samples and variants represented in the allele-frequency-filtered HPRC pangenome graph (AF- HPRC). SNVs and INDELs are shown separately across autosomes. Green indicates variants shared between the benchmark normal sample and the AF-HPRC graph, yellow indicates AF-HPRC-only variants, blue indicates COLO829BL-only variants, and pink indicates HG008N-only variants. (f) SBS96 mutational spectra and COSMIC signature decomposition for COLO829T and HG008T. COLO829T is dominated by UV-associated mutational signatures, whereas HG008T shows a more heterogeneous composition including age-related, APOBEC-associated, polymerase-related, and treatment-associated signatures.

Once a candidate position is identified within a node, Pansoma uses that position as the center anchor for tensor construction (Fig. 2b). The tensor contains five complementary channels: observed read bases, base quality, mapping quality, mismatch flag, and CIGAR operation. Nucleotide bases and alignment features, including substitutions, insertions, and deletions, are numerically encoded, allowing the model to learn local evidence patterns that distinguish true somatic variants from sequencing errors, germline variation, and graph-alignment artifacts (Fig. 2c). By combining population-aware graph alignment with node-based tensor construction, Pansoma transforms conventional somatic variant calling into a graph-native learning problem.

### Benchmark datasets

To support both model development and evaluation, we evaluated Pansoma in tumor-only mode using three benchmark resources with distinct biological and technical properties, each representing a complementary somatic variant-calling scenario. The SMaHT COLO829 benchmark [22] is derived from the COLO829 melanoma tumor cell line and its matched normal sample, COLO829BL, providing a well-characterized tumor-normal system for evaluating somatic SNV and indel detection across simulated tumor purities and sequencing technologies. The GIAB HG008 benchmark [26], developed through the Cancer Genome in a Bottle effort, includes multi-platform whole-genome sequencing data from the HG008-T pancreatic tumor and matched normal HG008 sample, enabling evaluation in an independent tumor type and testing cross-dataset generalizability. We also included the SMaHT HapMap mixture benchmark [22], in which predefined mixtures of multiple HapMap cell lines enable evaluation of low-VAF variant detection near the lower allele-fraction limit. Together, these datasets allowed us to benchmark Pansoma across natural tumor-derived datasets and a challenging synthetic low-VAF setting. For benchmark evaluation, we applied all tools in a tumor-only setting, without using matched normal samples as controls. For COLO829T and HG008T, Pansoma was compared with the tumor-only callers ClairS-tumor-only (ClairS-TO) and DeepSomatic-tumor-only (DeepSomatic-TO) across WGS, PacBio and ONT sequencing data, whereas for the HapMap benchmark, a specialized low-VAF Pansoma model was compared with Mutect2.

These datasets span markedly different variant allele fraction (VAF) regimes (Fig. 2d). COLO829T and HG008T contain variants distributed across a broad VAF range in both Illumina and PacBio data, with SNVs concentrated around intermediate allele fractions and INDELs showing broader, more platform-dependent distributions. In contrast, the HapMap benchmark is dominated by very low-VAF variants, reflecting its intended mixture design. In the HapMap mixture, HG005 makes up the majority background component at 83.5%, while five minor HapMap components contribute lower-frequency alleles: HG02622 at 10%, HG02486 at 2%, HG02257 at 2%, HG002 at 2%, and HG00438 at 0.5%. Thus, germline variants from the minor mixture components serve as controlled proxies for low-frequency somatic-like mutations.

The benchmark datasets also capture biologically distinct mutational processes. SBS96 mutational spectra and COSMIC [28] signature decomposition showed that COLO829T is dominated by UV-associated signatures, consistent with its melanoma origin (Fig. 2f). In contrast, HG008T displayed a more heterogeneous signature composition, including age-related, APOBEC-associated, polymerase-related, and treatment-associated components. These differences are important for benchmarking because they test whether Pansoma preserves biologically plausible somatic signal across tumors with distinct mutational processes rather than only optimizing aggregate variant-level metrics.

We next asked whether the pangenome graph could provide a population-aware germline prior for tumor-only somatic variant calling. Germline variants from COLO829BL and HG008N were compared with variants deconstructed from the allele-frequency-filtered HPRC graph (AF-HPRC; Fig. 2e). We used Dipcall [29] to identify germline variants from the benchmark normal samples and vg deconstruct [30] to extract graph-encoded variants relative to the GRCh38 path from the AF-HPRC graph. Across autosomes, more than 79% of normal-sample SNVs and 73% of INDELs overlapped variants represented in the AF-HPRC graph, shown by the green bars in Fig.2e. This overlap demonstrates that many inherited alleles present in the benchmark individuals are already encoded as graph paths. For tumor-only analysis, this representation is important because reads carrying these germline alleles can align to the corresponding graph haplotypes instead of being forced onto a single linear reference, where they may appear as mismatches, indels, or local alignment artifacts.

### Pansoma performance evaluation

We evaluated Pansoma in tumor-only mode across three complementary benchmark settings designed to test different aspects of somatic variant calling, with the technique details described in Methods. For COLO829T and HG008T, Pansoma was compared with ClairS-TO and DeepSomatic-TO across WGS, PacBio and ONT sequencing data. For the HapMap benchmark, a specialized low-VAF Pansoma model was compared with Mutect2. The detailed results are provided in Supplementary Table 2, 3.

Across COLO829T and HG008T, Pansoma achieved the highest SNV F1-scores in nearly all platform and purity settings (Fig. 3a, b). In the COLO829T benchmark, Pansoma maintained strong SNV performance across simulated tumor purities from 25% to 100%, indicating robustness across a broad range of tumor-content settings. In the independent HG008T benchmark, Pansoma also achieved the highest SNV F1-scores across WGS, PacBio and ONT datasets, supporting its generalizability across tumor type and sequencing platform. Precision and recall analyses showed that these F1-score gains were largely driven by improved precision (Supplementary Fig. 1). These performance gains likely reflect the ability of Pansoma to reduce the background candidate burden that is intrinsic to tumor-only somatic variant calling. We further investigate the false-negative calls (Supplementary Figs. 2 and 3) and find that many are associated with graph alignment behavior. Some false-negative variants overlap existing nodes in the pangenome reference, suggesting that a subset of the ground-truth somatic variants may correspond to germline alleles already represented in the pangenome. Without a matched normal control, true somatic variants must be distinguished from a much larger background of germline alleles, sequencing artifacts and alignment-induced candidates. By using population-aware pangenome graph alignment, Pansoma can reduce apparent mismatches and gap events caused by forcing sample-specific alleles onto a single linear reference, while graph-derived pileup tensors allow the model to learn local somatic-variant features directly from graph-aligned reads.

**Figure 3.**
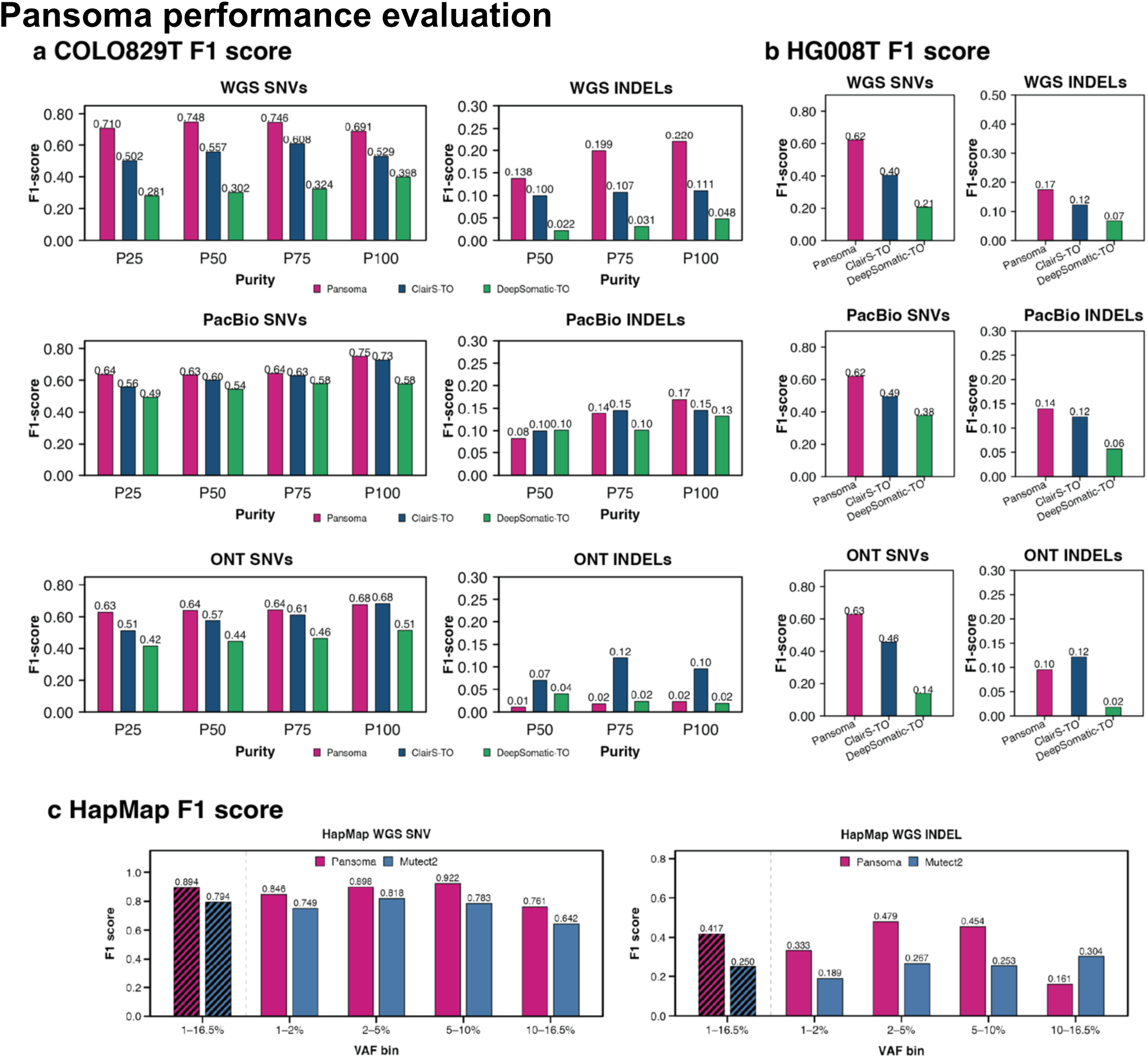
Performance evaluation of Pansoma in tumor-only somatic variant calling across benchmark datasets (a) F1-score comparison of Pansoma, ClairS-TO and DeepSomatic-TO on the COLO829T benchmark across WGS, PacBio and ONT datasets. Results are shown separately for SNVs and INDELs across different tumor purity levels. Pansoma achieved the highest SNV F1-scores across most purity levels and sequencing platforms, while INDEL performance was more platform-dependent. (b) F1-score comparison of Pansoma, ClairS-TO and DeepSomatic-TO on the HG008T benchmark across WGS, PacBio and ONT datasets. SNV and INDEL results are shown separately for each sequencing platform. Pansoma showed consistently strong SNV performance and achieved the highest INDEL F1-scores in WGS and PacBio, whereas ClairS-TO performed better for ONT INDELs. (c) Evaluation of Pansoma on the SMaHT HapMap WGS benchmark across variant allele fraction (VAF) bins. A specialized low-VAF Pansoma model was compared with Mutect2 for SNV and INDEL detection. Hatched bars summarize performance across the combined 1–16.5% VAF interval, while the remaining bars show performance within individual VAF bins.

We compared graph-based and GRCh38-based alignments across COLO829T and HG008T. Graph alignment consistently increased the number of perfectly aligned reads while reducing alignment discrepancies relative to linear-reference alignment. Across both datasets, graph alignment increased perfect alignments by up to 11% in WGS data and by more than four-fold in long-read datasets, while substantially reducing soft-clipped bases, substitutions and indel events (Supplementary Fig. 4). These results indicate that population-aware graph references better represent sample-specific haplotypes and reduce alignment artifacts that arise when reads are forced onto a single linear reference.

Consistent with these alignment improvements, graph-based candidate generation substantially reduced the number of putative somatic-variant candidates. Using identical thresholds of more than two variant supporting reads and variant allele frequency greater than 10%, graph-based candidate generation produced fewer candidate records than GRCh38-based candidate generation across chromosomes 1–22 in both COLO829T and HG008T on Illumina data. The graph alignment reduced SNV candidate examples to 63.8% and INDEL candidate examples to 24.5% of the corresponding GRCh38-based counts, resulting in an overall reduction of candidate records to less than 64% of those generated from the linear reference (Supplementary Fig. 5). By reducing alignment-induced mismatches and gap events before machine-learning classification, graph alignment decreases the background candidate burden that must be distinguished from true somatic variants, providing a likely explanation for the improved precision and F1-scores observed for Pansoma.

In the HapMap low-VAF benchmark, the specialized Pansoma model outperformed Mutect2 across most SNV VAF bins and improved INDEL detection in the 1–10% VAF range (Fig. 3c). ClairS-TO and DeepSomatic-TO were not included in this low-VAF comparison because their available tumor-only workflows were not designed for ultra-deep tumor-only somatic variant calling below 10% allele fraction in this benchmark setting. Pansoma showed lower performance in the 10–16.5% INDEL bin, where Mutect2 achieved a higher F1-score. This pattern may reflect the VAF distribution of the HapMap mixture, in which most somatic-like variants are concentrated below 10% VAF (Fig. 2d). As a result, higher-VAF bins contain fewer true positives while retaining background artifacts, making F1-scores in these bins more sensitive to data imbalance and false-positive calls. Overall, these results demonstrate that the low-VAF Pansoma model improves tumor-only somatic variant detection in ultra-deep sequencing data, particularly for SNVs and low-to-moderate VAF INDELs.

### Graph-aware analysis of Pansoma SNV predictions

To better understand Pansoma beyond aggregate F1-scores, we performed graph-aware analyses of SNV predictions across caller overlap, graph-coordinate representation and PoN-independent performance. These analyses addressed three questions: whether Pansoma predictions retain biologically plausible tumor mutational signal, whether non-reference graph paths provide additional variant evidence beyond the GRCh38 reference path, and whether pangenome alignment and representation can reduce background calls without panel-of-normals filtering.

We first compared SNV predictions from Pansoma, ClairS-TO and DeepSomatic-TO with the benchmark truth sets across HG008T and COLO829T WGS, PacBio and ONT datasets (Fig. 4a). Pansoma recovered a shared core of truth-overlapping calls across sequencing platforms, while each method also produced caller-specific SNV sets not shared by the others. Because the total number of calls differed substantially among methods, the Venn diagrams are intended to illustrate the structure of overlap rather than to provide area-scaled quantitative summaries; detailed call counts are provided in Supplementary Table 4, 5. We next examined whether these caller- associated SNV sets retained recognizable tumor mutational signal by analyzing their COSMIC mutational signature compositions. In COLO829T, Pansoma predictions showed clear contributions from UV-associated signatures, consistent with the melanoma origin of COLO829T and the benchmark mutational profile. In contrast, HG008T showed a more heterogeneous signature composition, consistent with its more complex mutational landscape and the absence of a single dominant mutational process. To quantify the similarity between caller-specific and core- overlapping signatures, we represented each mutational signature profile as a vector and calculated the cosine similarity between each caller and the benchmark ground-truth profile, with the results shown as bar plots in Fig. 4a. We also calculated the cosine similarity between the COLO829T graph-coordinate somatic variant signature profiles identified by Pansoma and the benchmark profile in Fig. 4c. Additionally, the lower cosine similarity observed for HG008T may be due to the more complex benchmark signature profile, which is distributed across several major components. By contrast, COLO829T has a more concentrated signature profile dominated by UV-associated signatures, allowing the predicted profiles to more closely match the benchmark.

**Figure 4.**
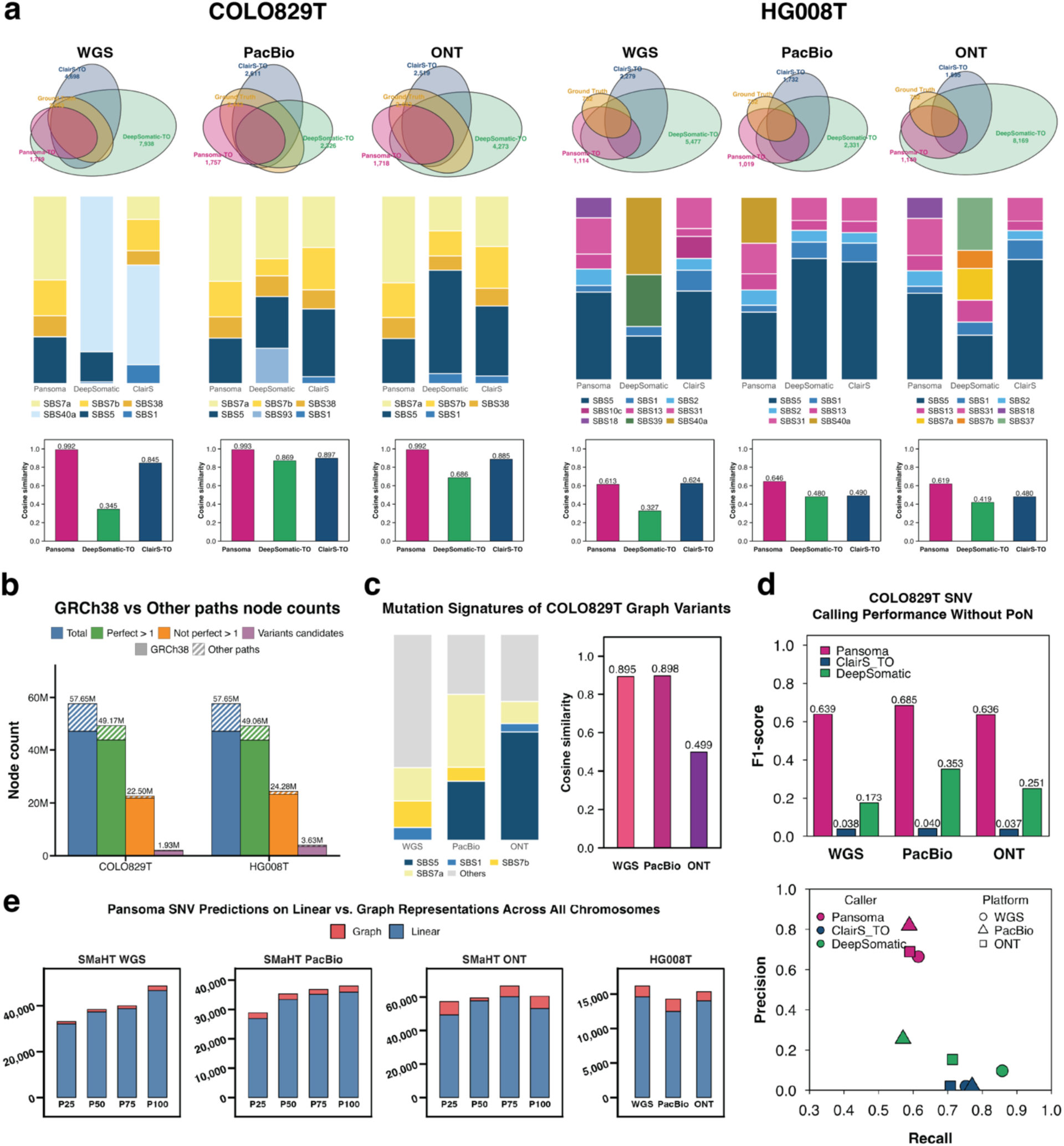
Caller-specific SNV predictions, graph-coordinate variant representation and PoN- independent performance. (a) Overlap of PASS SNV calls among Pansoma, DeepSomatic-TO, ClairS-TO and the benchmark truth set for HG008T and COLO829T across WGS, PacBio and ONT datasets. Venn diagrams show shared and caller-associated SNV sets, and stacked bar plots below summarize the COSMIC mutational signature composition of SNVs from each caller, and the bar plots below show the cosine similarity between each caller’s mutational signature profile and that of the benchmark truth set. (b) Distribution of graph nodes and candidate-containing nodes across the GRCh38 reference path and non-reference graph paths for HG008T and COLO829T across all autosomes. Bars summarize total graph nodes, read-supported nodes, nodes with non-perfect read alignments and variant candidate nodes. (c) COSMIC mutational signature composition of COLO829T graph-coordinate SNV predictions retained outside the linear-reference representation across WGS, PacBio and ONT datasets, along with the cosine similarity. (d) COLO829T SNV calling performance without panel-of-normals filtering. The precision–recall plot compares Pansoma, ClairS-TO and DeepSomatic-TO across WGS, PacBio and ONT datasets, and the bar plot shows the corresponding F1-scores. (e) Composition of Pansoma SNV predictions across linear-reference and graph-coordinate representations. Stacked bars show the number of SNV predictions projected to the linear reference and retained as graph-coordinate variants across SMaHT WGS, PacBio, ONT and HG008T datasets.

We next asked whether the pangenome graph provided additional candidate space beyond the GRCh38 reference path. Across all autosomes, a substantial fraction of graph nodes belonged to non-reference paths and reads from both HG008T and COLO829T supported many of these nodes (Fig. 4b). Among read-supported nodes, nodes with non-perfect read alignments, including mismatches or gap-related events, represented regions with potential variant evidence. A smaller subset of these nodes passed Pansoma’s candidate-generation criteria based on allele fraction and read support. Importantly, candidate-containing nodes were not restricted to the GRCh38 path but also occurred on non-reference graph paths, indicating that graph alignment exposes candidate variation that is not naturally represented in a single linear-reference coordinate system.

This graph-derived signal was also retained in the final prediction outputs. Across SMaHT WGS, PacBio, ONT and HG008T datasets, most Pansoma SNV predictions could be projected back to the linear reference, but a considerable fraction remained represented only in graph coordinates (Fig. 4e). Notably, ONT datasets showed a higher proportion of graph-coordinate predictions than WGS or PacBio datasets. This pattern may reflect the higher error rate and more complex alignment profiles of ONT reads, which can generate more candidate variants on alternative graph paths and increase the fraction of predictions retained outside the linear-reference representation. Thus, the contribution of the graph is not limited to intermediate alignment or candidate generation; it also expands the final representation by preserving predictions from non-reference graph paths. These graph-coordinate calls provide an additional layer of tumor-supported variant evidence that would be lost or difficult to express in conventional linear-reference VCF outputs.

To evaluate whether graph-coordinate predictions retained biologically relevant signal, we further analyzed the mutational signatures of COLO829T SNV predictions retained outside the linear-reference representation (Fig. 4c), and we also performed the same analysis for HG008T and present the results in Supplementary Fig. 7. These graph-coordinate predictions contained COLO829-relevant signature components, including UV-associated signatures. Although graph- coordinate calls remain difficult to benchmark directly with current linear-reference truth sets, this signature pattern indicates that they retain tumor-associated mutational signal rather than appearing as an unstructured technical byproduct of graph alignment. We focused this signature analysis on COLO829T because its UV-dominated mutational profile provides a more distinctive biological signal for evaluating graph-coordinate predictions, whereas HG008T shows a more heterogeneous signature composition.

We then evaluated whether graph-based alignment and representation reduced the dependence of tumor-only calling on external background filters. In conventional tumor-only workflows, panel-of-normals filtering is commonly used as an empirical prior to suppress recurrent sequencing artifacts, germline-like variants and other non-somatic background signals. By removing PoN filtering, we created a setting in which caller performance depends more directly on read alignment, genome representation and model inference. In this PoN-independent setting, Pansoma maintained substantially higher precision and F1-scores than ClairS-TO and DeepSomatic-TO across WGS, PacBio and ONT datasets (Fig. 4d). These results suggest that pangenome alignment provides population- and haplotype-aware context that helps suppress background false positives in tumor-only somatic variant calling, even without relying on a separate PoN resource.

As an exemplar read-level illustration of graph-coordinate variant representation, we examined a representative SNV located within an alternative sequence segment represented in HG008N haplotype 1, the haplotype-resolved assembly of the matched normal sample provided with the HG008T benchmark [27], and HPRC v1.1 AF-Graph node 14673857, but not directly on the GRCh38 reference path (Fig. 5). Compared with the GRCh38 view, the haplotype and graph-node views provide a more contiguous local sequence context with read support at the candidate position. This IGV screenshot [31] example illustrates how graph-coordinate reporting can preserve variant evidence on non-reference haplotype paths that is difficult to express or interpret using conventional linear-reference coordinates.

**Figure 5.**
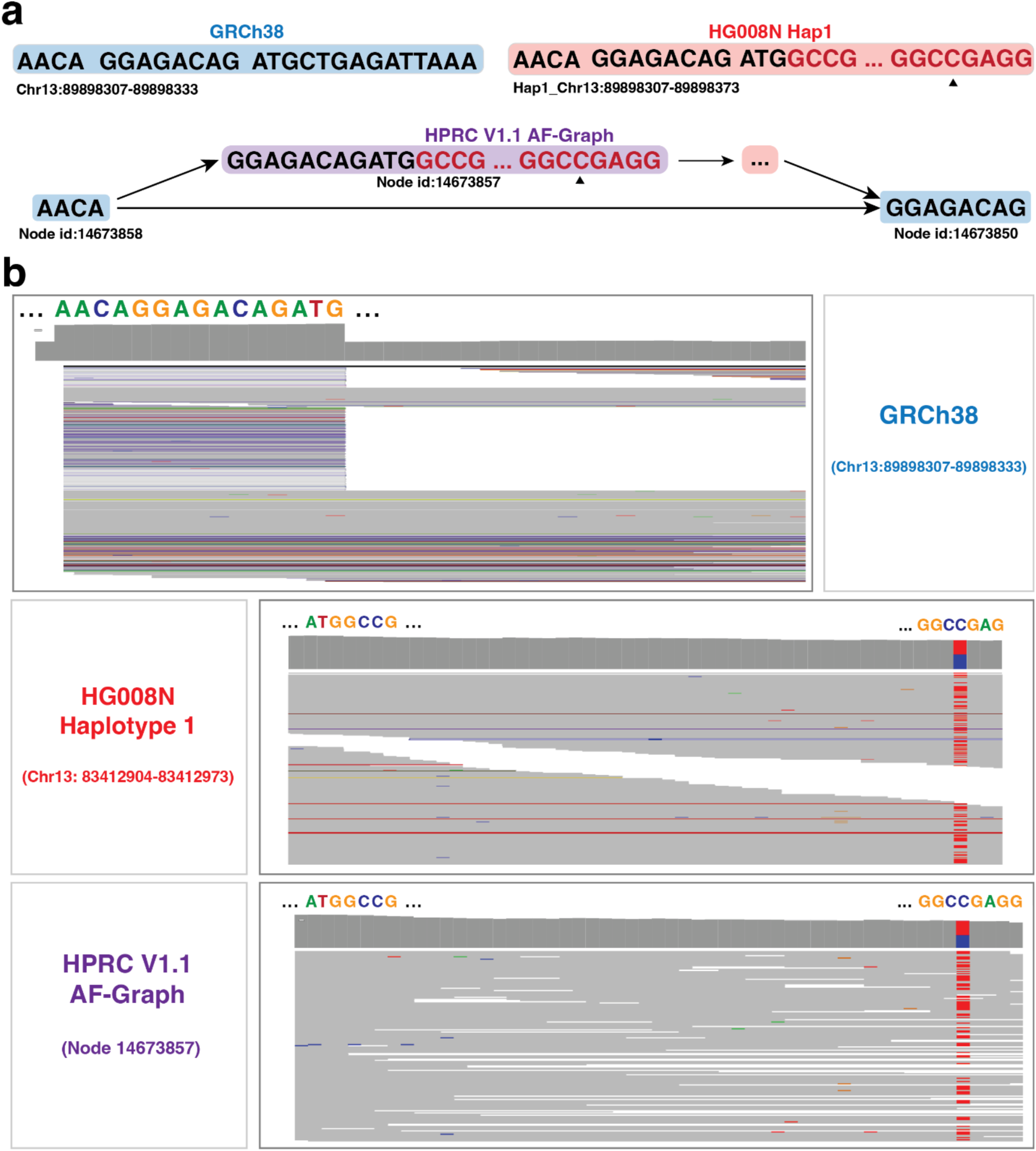
Representative graph-coordinate SNV supported by haplotype and graph-aligned read pile up. (a) Local sequence context of a graph-coordinate SNV on chromosome 13. The candidate SNV is located within an alternative sequence segment represented in HG008N haplotype 1 and HPRC v1.1 AF-Graph node 14673857 but not directly represented on the GRCh38 reference path. The SNV position is highlighted in red. (b) Read-level pileups for the corresponding GRCh38 locus, HG008N haplotype 1 sequence and HPRC AF-Graph node. The haplotype and graph views provide a contiguous alternative representation with read support for the highlighted SNV. Screenshots were generated using the IGV-Web browser.

Together, these aggregate and read-level analyses indicate that Pansoma not only improves tumor-only SNV prediction but also provides a graph-aware representation of somatic variant evidence. By integrating pangenome-based alignment, graph-coordinate candidate discovery and graph-specific variant reporting, Pansoma preserves tumor-associated variant signals on graph paths that are difficult to access through conventional linear-reference workflows.

## Methods

### Node pileup generation and graph alignment

One key challenge in performing variant calling on graph alignments is it differs fundamentally from conventional linear-reference alignments. Standard graph alignment formats such as GAM and GAF preserve read-to-graph alignment information, but they are not optimized for high-throughput random access by graph node. To call variants within a graph region, existing approaches typically require traversing the GAM or GAF file to collect supporting reads or loading the full alignment file into memory. This makes direct pileup generation from graph alignment files inefficient for genome-wide somatic variant calling.

To address this limitation, we developed a node pileup format, referred to as NPU, to store graph alignment in a node-centric manner and support random access with the node ID index file. In summary, reads are organized into blocks according to the graph node IDs they align with, while the accompanying index file stores the physical offsets for each node-specific block. This design enables the NPU format to rapidly retrieve all reads aligned to a queried node ID without scanning the entire alignment file. The detailed layout of the NPU data and index structures is summarized in Supplementary Table 1.

In this work, sequencing reads were aligned to the human pangenome graph using vg giraffe [18] version 1.66.0 to generate graph alignment files. For long-read sequencing data, additional vg giraffe presets were applied: “-b hifi” for PacBio HiFi data and “-b r10” for ONT data.

### Tensor construction and model training

Candidate variant sites were identified from node-level pileups at single-base resolution. For each position within a retained graph node, Pansoma first excluded reads with mapping quality ≤ 5, then summarized the observed read alleles and compared them with the graph reference sequence. A position was retained as a candidate if the non-reference allele frequency exceeded 10% and it was supported by more than two reads. This hard-filtering step reduced the search space from all graph positions to a smaller set of putative variant sites.

For each candidate variant site, Pansoma converts the corresponding graph-node pileup into a multi-channel tensor suitable for analysis by a convolutional neural network. A fixed-width local window is extracted around the candidate site, with the candidate position placed at the center of the window. In our implementation, the tensor width was set to 101 bp, and the maximum read depth was set to 200 reads. If the candidate-centered window extended beyond the boundary of the graph node, the window was truncated to the available node sequence.

Each candidate pileup was represented as a three-dimensional tensor with dimensions C × W × H, where C is the number of feature channels, W is the width of the local genomic window, and H is the maximum read depth. The horizontal axis of the tensor represents positions in the candidate-centered window, whereas the vertical axis represents reads overlapping the candidate region. Reads covering the window were sorted by their assigned haplotype path and start position to create a structured image-like representation. If the number of overlapping reads was lower than the maximum depth, the remaining rows were padded with zeros. If the read depth exceeded the maximum depth, reads were subsampled while preserving the observed allele-frequency structure at the candidate site.

Each tensor contained five feature channels to build the image-like tensors.

1. **Read Bases:** The nucleotide sequence of each read fragment within the window. Each position can be (A, C, G, T, N), where N stands for null.
2. **Base Quality:** The Phred-scaled base quality score for each nucleotide in the read, normalized to a range of [0, 1]. This channel provides the model with confidence information for each base call.
3. **Mapping Quality:** The mapping quality (MAPQ) score of each read alignment is also normalized to [0, 1]. This value is constant across all bases for a single read and indicates the confidence that the read is mapped to the correct location in the pangenome graph.
4. **Mismatch flag:** Indicates agreement between each read base and the reference: 0 represents a match, 1 a mismatch, and −1 padding. At the variant anchor, a mismatch is amplified to 5, emphasizing the focal candidate position for the model.
5. **CIGAR operation:** This channel records the local alignment operation for each read within the candidate window, including match/mismatch, insertion, deletion, or clipping states. By encoding these CIGAR-derived patterns, the model can learn whether the observed read evidence reflects a potential variant signal or an alignment/sequencing artifact.

Pansoma used a convolutional neural network to classify candidate tensors as somatic or non-somatic (Supplementary Fig. 6). The network begins with a convolutional stem that reduces spatial dimensions and increases feature depth, followed by four stages of ConvNeXt-style convolutional blocks with LayerNorm and GELU activation. The four stages use depths of [3, 3, 27, 3] and widths of [192, 384, 768, 1536]. Within each block, depth-wise convolutions capture local sequence and alignment patterns, while the multilayer perceptron expands and contracts feature dimensions. Channel and spatial attention mechanisms were incorporated through CBAM to highlight informative features, and gated residual normalization was used to stabilize training. Residual connections with stochastic depth were applied throughout the network, and downsampling layers progressively reduced spatial resolution while increasing feature capacity. The final representation was aggregated by global average pooling and passed to a fully connected classifier.

For each sequencing platform, candidates from chromosomes 2–22 were used for model development and divided into training and validation subsets using a fixed random seed. The validation set was used only for checkpoint selection, whereas chromosome 1 was held out for final testing. HG008T was excluded from model fitting and served as an independent external test dataset. This design prevented overlap of genomic loci between model development and testing and enabled evaluation across samples.

Model training was performed using PyTorch Distributed Data Parallel on four NVIDIA H100 GPUs. The model was optimized with AdamW [32] using an initial learning rate of 1 × 10⁻^⁴^ and a weight decay of 0.01. Training used 3 epochs of linear warm-up followed by cosine annealing for a total of 80 epochs, with both learning rate and weight decay annealed over training. To address class imbalance between true somatic variants and negative candidates, we used class-weighted cross-entropy loss with a positive-class weight of 50 for WGS and PacBio data, but 200 for ONT data. Model checkpoints were saved beginning at epoch 5 and subsequently whenever the validation F1-score of the positive class improved.

### Projection between linear and graph coordinates and VCF generation

After model inference, Pansoma predictions were initially represented in graph coordinates, including graph node ID, node offset, reference allele, alternate allele, and model probability. To generate standard VCF output, predicted variants were classified according to whether their graph locations could be projected back to the GRCh38 reference path. Variants located on reference- path nodes or on alternative nodes that could be unambiguously projected to GRCh38, were converted into standard linear-reference VCF records. Variants located on alternative graph paths without an unambiguous GRCh38 projection were retained as graph-coordinate VCF records.

A coordinate mapping was constructed between GRCh38 reference positions and graph coordinates, enabling benchmark variants to be assigned to graph node IDs and node offsets whenever corresponding graph positions were available. To build this mapping, we first collected all graph nodes traversed by the GRCh38 path in the NPU representation and identified the corresponding GRCh38 loci associated with these nodes. For nodes located on alternative haplotype paths, we additionally collected the adjacent 1-bp GRCh38-linked SNV nodes and used their related GRCh38 positions to infer the coordinates of graph-only positions. In this case, we get this coordinate mapping as a hash table, which allowed Pansoma to use existing linear-reference truth sets for model development while preserving graph-specific predictions that cannot be represented directly in linear coordinates.

### Panel-of-normals filtering

After linear-reference VCF generation, panel-of-normals filtering was applied to reduce residual germline and recurrent background calls. In this work, we apply four different panels of normals (PoNs) [15, 33–35] to further filter the linear-reference variants. PoN 1 was derived from the Broad/GATK af-only gnomAD hg38 resource [33], which is a stripped-down gnomAD allele- frequency resource commonly used as a germline background set. PoN 2 was based on dbSNP build 138 hg38 and was restricted to non-somatic sites [15]. PoN 3 was the Broad/GATK 1000G PoN hg38, an official panel of normals generated from 1000 Genomes Project samples [34]. PoN 4 was derived from CoLoRSdb v1.1.0, a long-read population variant database [35], from which sites with AF ≥ 0.001 were retained.

These resources were selected to capture complementary classes of common population polymorphisms, known non-somatic loci, recurrent background variation, and long-read- supported population variants. The PoN filtering step was applied only to variants represented in linear-reference coordinates, because these resources are defined on GRCh38 rather than graph coordinates.

### Benchmarking design and performance evaluation

We evaluated Pansoma in tumor-only mode using three benchmark resources: COLO829T, HG008T, and the SMaHT HapMap mixture datasets. The COLO829 benchmark included PCR- free Illumina whole-genome sequencing, PacBio Revio HiFi Fiber-seq data, and Oxford Nanopore Technologies long-read sequencing data. To assess performance across tumor purity levels, COLO829T tumor reads were combined with matched COLO829BL normal reads to generate in silico tumor-purity settings of 25%, 50%, 75%, and 100%. For each mixture dataset, graph alignments were generated, indexed node pileups were constructed, candidate variants were identified, and candidate tensors were generated for Pansoma inference.

For the HG008 benchmark, we used tumor sequencing data from HG008-T, a pancreatic ductal adenocarcinoma tumor sample, across three sequencing platforms. The short-read dataset was generated on the Illumina NovaSeq 6000 platform using PCR-free paired-end whole-genome sequencing. The long-read datasets included PacBio Revio HiFi reads and Oxford Nanopore ultra- long reads generated on the PromethION platform. Because only one tumor-purity setting was available for HG008T, this benchmark was used as an independent evaluation across WGS, PacBio, and ONT sequencing platforms.

For the HapMap benchmark, we used the WGS dataset from the SMaHT HapMap mixture, a synthetic admixture of six HapMap cell lines designed to generate controlled low-VAF somatic- like variants. In this dataset, HG005 makes up the majority background component at 83.5%, while the remaining cell lines contribute lower-frequency alleles: HG02622 at 10%, HG02486 at 2%, HG02257 at 2%, HG002 at 2%, and HG00438 at 0.5%. Because low-VAF detection requires high read depth, this analysis was limited to short-read WGS data. For this benchmark, we applied a specialized low-VAF Pansoma model: candidate variants with allele frequency greater than 10% were excluded, candidate variants with allele frequency greater than 1% were retained, and the maximum pileup depth was increased to 500 reads to accommodate the high sequencing depth of the mixture dataset.

For COLO829T and HG008T, Pansoma was compared with tumor-only configurations of ClairS- TO and DeepSomatic-TO. For the HapMap low-VAF benchmark, Pansoma was compared with Mutect2. All methods were evaluated from the same raw sequencing reads for each benchmark. Linear-reference callers were run using their standard GRCh38-based alignment and calling workflows, whereas Pansoma used vg graph alignment to the allele-frequency-filtered HPRC pangenome graph followed by graph-native candidate generation, tensor construction, and model inference.

Final Pansoma predictions were reconstructed from graph coordinates. Predictions that could be projected to GRCh38 were converted into conventional linear-reference VCF records and used for direct comparison with linear-reference truth sets and comparator call sets. Predictions that could not be unambiguously projected to GRCh38 were retained as graph-coordinate VCF records and analyzed separately. For performance evaluation, projected linear-reference calls were compared with benchmark truth variants using standard TP, FP, and FN classification. Precision was calculated as *TP*/(*TP* + *FP*), recall as *TP*/(*TP* + *FN*), and F1-score as 2 × *precision* × *recall*/(*precision* + *recall*). SNVs and INDELs were evaluated separately.

## Discussion

In this study, we developed Pansoma, a machine learning–based somatic variant caller that directly operates on the pangenome graph alignment. Compared with existing methods such as DeepSomatic and ClairS, Pansoma achieves robust precision and F1 performance in tumor-only settings across both short- and long-read sequencing data. This improvement can be credited to its graph-native design: by aligning reads to paths that encode common germline variants, Pansoma reduces false positives that typically arise when germline variation is misclassified as somatic in linear reference–based approaches.

A central innovation of Pansoma is its node pileup (NPU) representation and tensorization strategy, which transforms graph-based alignments into structured, image-like inputs for deep learning. This framework provides a new perspective for variant calling on graph references. While we primarily benchmarked linearized outputs, about 2% of variants located on alternative paths remain in the node-based outputs, which are difficult to benchmark. Importantly, the linear projection is not limited to GRCh38 and can be extended to CHM13 or other references, enabling interoperability between graph-native and conventional workflows. Meanwhile, the node-based VCFs generated by Pansoma provide a foundation for deeper exploration as graph-based tools continue to mature.

As the human pangenome expands to incorporate more haplotypes and structural variants, we anticipate further gains in accuracy and generalizability. Beyond serving as an alternative to reference-based pipelines, graph-native variant calling provides a unifying framework to connect assembly-based and population-scale variation into a single representation. Together, these advances highlight the potential of pangenome-native AI methods to transform somatic variant detection and accelerate the next generation of precision genomics.

Several limitations remain. First, graph-coordinate somatic variants are difficult to benchmark because current truth sets are primarily defined on linear references. Second, candidate generation thresholds may limit sensitivity for variants below the standard VAF cutoff, although a specialized low-VAF model partly addresses this limitation. Third, INDEL calling, especially in ONT data, remains challenging and platform dependent. Fourth, performance depends on the composition and filtering of the pangenome graph, and sample-relevant haplotypes may influence alignment and candidate representation. Finally, downstream annotation and clinical interpretation tools remain largely linear-reference based, requiring projection steps which cause loss of some graph-specific information.

## Data availability

The COLO829T, COLO829BL and HapMap mixture sequencing datasets and benchmark resources used in this study are available through the Somatic Mosaicism across Human Tissues (SMaHT) Network. The HG008T and HG008N sequencing datasets and somatic small-variant benchmark are available through the Genome in a Bottle Consortium. The HPRC pangenome graph used in this study is publicly available through the Human Pangenome Reference Consortium. The pretrained Pansoma models are available through Zenodo at https://doi.org/10.5281/zenodo.21483622. Source data are provided with this paper.

## Code availability

The Pansoma source code, including programs for node pileup generation, tensor construction, model training and inference, and variant reconstruction, is publicly available at https://github.com/Jiawei-Shen/Pansoma. Scripts used to reproduce the benchmarking analyses, figures and tables are available in the same repository.

## Acknowledgements

We thank the Human Pangenome Reference Consortium and the Somatic Mosaicism across Human Tissues Network for generating and making available the pangenome resources, sequencing datasets and benchmark variant sets used in this study. We thank Justin Zook and colleagues in the Genome in a Bottle Consortium for providing early access to the HG008 sequencing data and somatic small-variant benchmark resources. We also thank the developers and maintainers of the open-source software and computational resources that supported this work.

