## Supplement for "Pansoma, a machine learning tool for identifying somatic variants using pangenome graphs"

### Supplementary

#### Supplement Figure 1

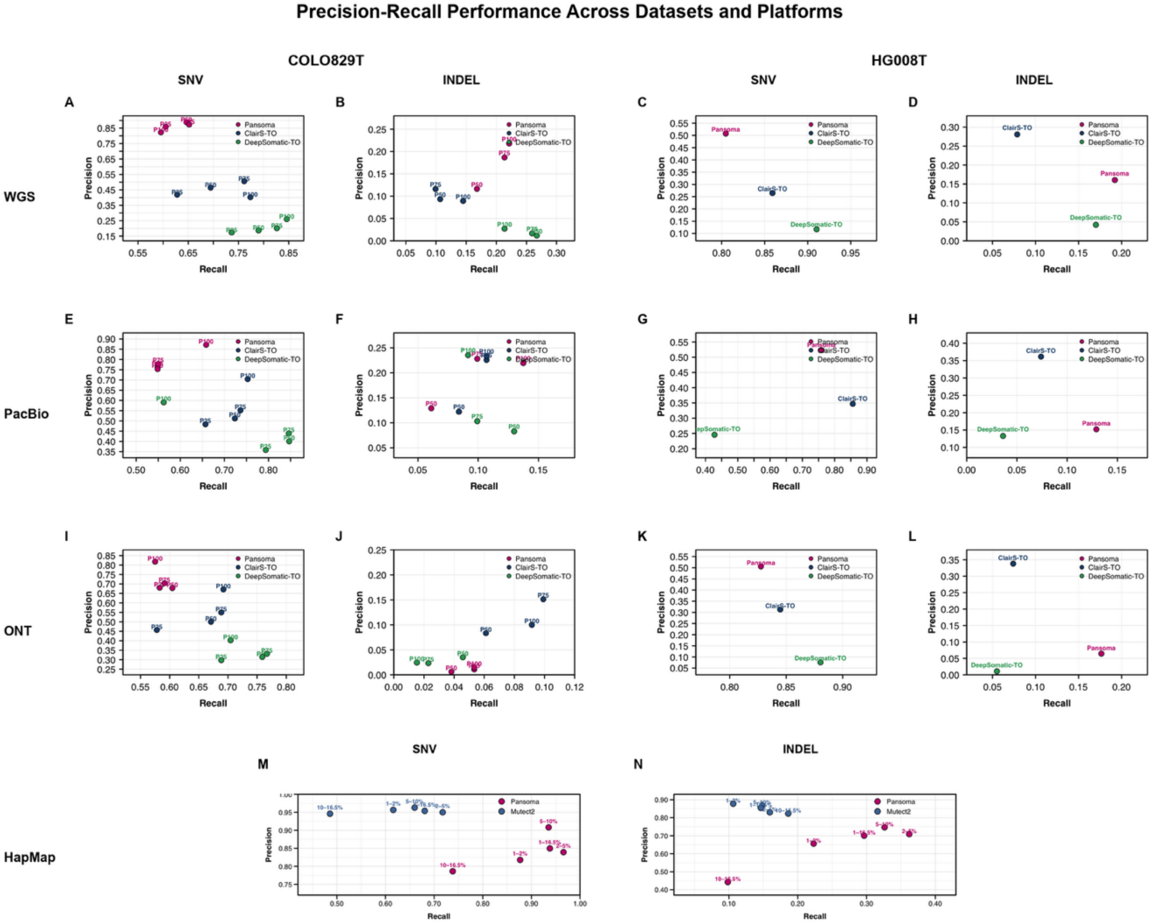


**Supplementary Figure 1. Precision–recall performance across datasets and sequencing platforms.**

Precision and recall of SNV and INDEL calls were compared across COLO829T and HG008T datasets for WGS, PacBio HiFi and ONT platforms. Panels A–L show performance of Pansoma, ClairS-TO and DeepSomatic-TO, with COLO829T points labeled by tumor purity level. Panels M–N show low-VAF HapMap WGS benchmark performance for SNVs and INDELs, comparing Pansoma with Mutect2 across VAF bins.

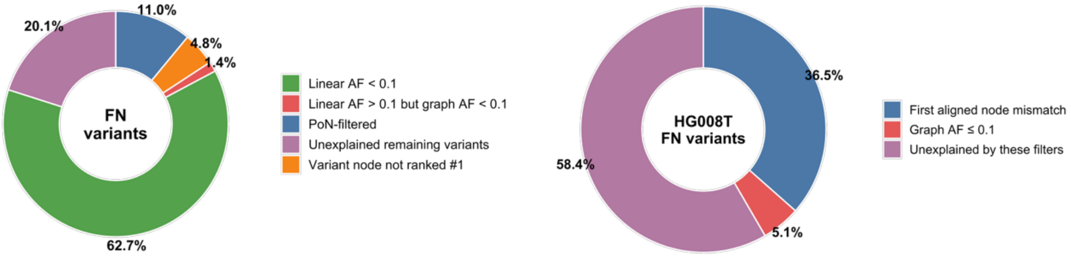


**Supplementary Figure 2. Stepwise classification of missed variants in graph-aware Pansoma analysis.**

Donut charts summarize the sequential breakdown of false-negative variants after applying graph- and linear-coordinate evidence. The HG008T analysis attributes missed variants to graph-alignment consistency and graph-support filters, including insufficient graph allele fraction or read support. The COLO829T analysis further separates missed variants by PoN filtering, non-dominant variant-node support, discordant linear versus graph allele fraction, low GRCh38 allele fraction and remaining unexplained variants. Percentages indicate the fraction of missed variants assigned to each stepwise category.

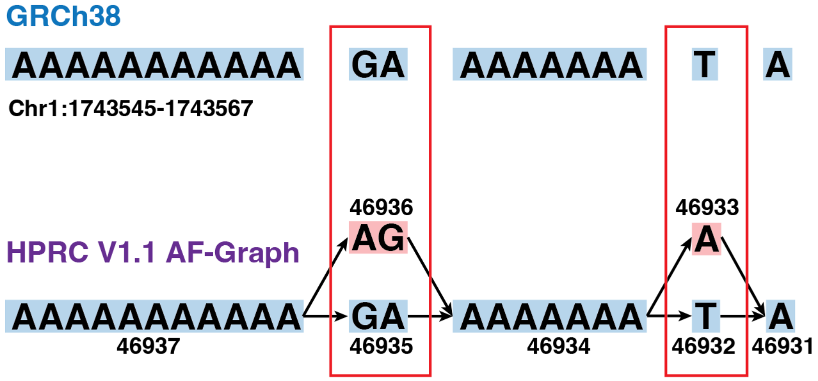


**Supplementary Figure 3. False negative examples.**

Representative HG008T false-negative examples caused by repetitive sequence context and GRCh38 reference bias. Three candidate SNVs, chr1:1,743,557 G>A, chr1:1,743,558 A>G, and chr1:1,743,566 T>A, appear as mismatches relative to the GRCh38 linear reference. In the HPRC v1.1 AF-Graph, however, the local haplotype sequence is represented by alternative graph paths, allowing reads to align perfectly without supporting the GRCh38-projected variant nodes. This graph-resolved alignment reduces the apparent allele frequency of these candidate sites to nearly zero, leading Pansoma not to call them as variants. These cases therefore reflect GRCh38 reference-bias artifacts rather than true missed variants.

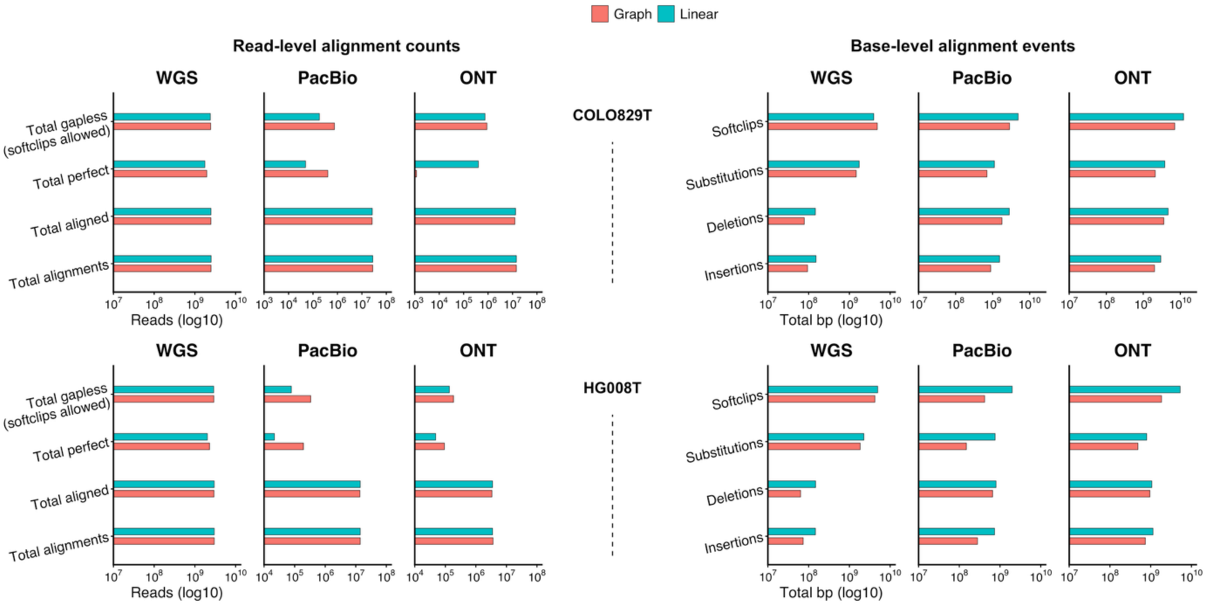


**Supplementary Figure 4. Alignment characteristics of graph and linear-reference mapping across sequencing platforms.**

Read-level alignment counts and base-level alignment events were compared between graph-based and GRCh38 linear alignments for COLO829T and HG008T across WGS, PacBio HiFi and ONT data. Left panels show total alignments, aligned reads, perfect alignments and gapless alignments, while right panels summarize soft-clipped bases, substitutions, deletions and insertions. All counts are shown on a log10 scale.

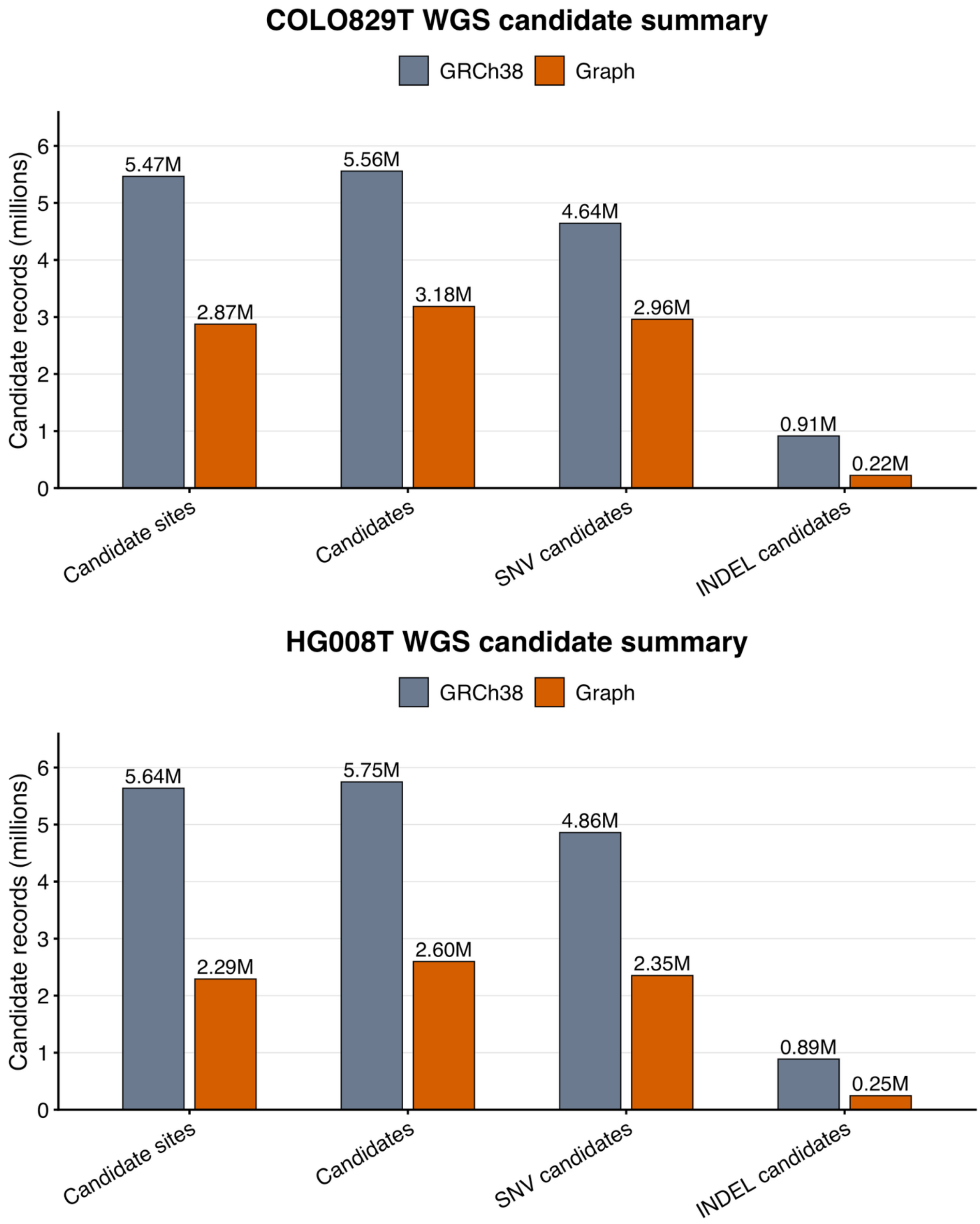


**Supplementary Figure 5. GRCh38 versus AF-HPRC graph candidate-generation comparison.**

Candidate-generation summaries for WGS datasets from COLO829T and HG008T using GRCh38-based linear alignment and AF-HPRC graph alignment. Bars show the number of candidate sites, total candidate records, SNV candidates and INDEL candidates generated using the same filtering thresholds. Compared with GRCh38, AF-HPRC graph-based candidate generation substantially reduced the total candidate burden in both datasets, with the strongest reduction observed for INDEL candidates.

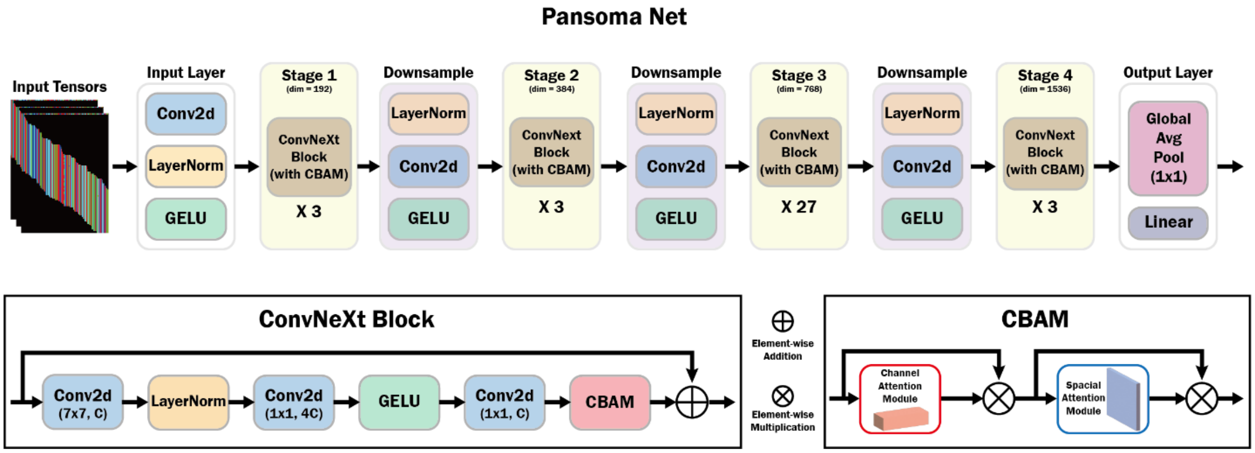


**Supplementary Figure 6. Pansoma neural network architecture.**
Pansoma uses a ConvNeXt-based classifier with CBAM attention to predict somatic variants from input pileup tensors. The network contains a convolutional stem, four ConvNeXt stages with dimensions of 192, 384, 768 and 1536, inter-stage downsampling layers, global average pooling and a linear classification head. Each ConvNeXt block includes depthwise convolution, layer normalization, pointwise convolutions, GELU activation, CBAM attention and a residual connection. The CBAM module refines features using sequential channel and spatial attention.

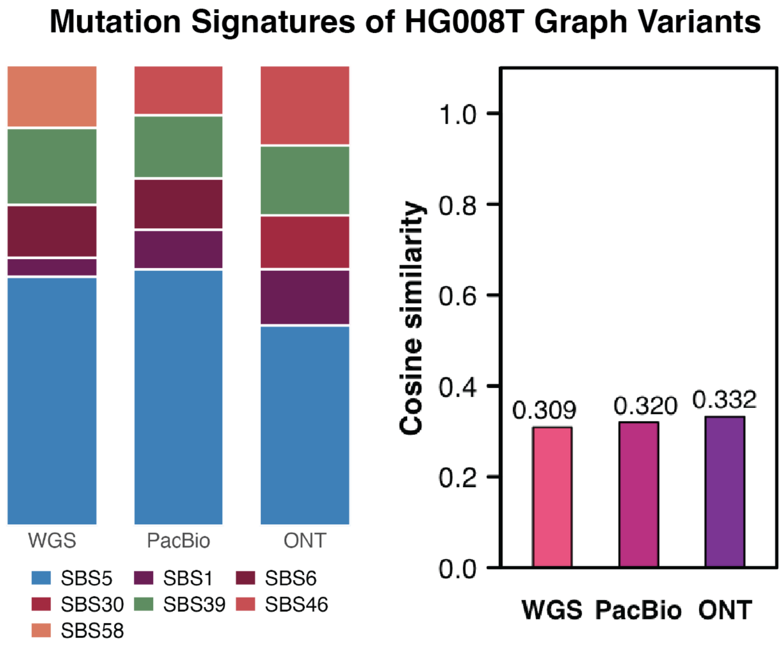

**Supplementary Figure 7. HG008T graph-coordinate SNV mutation signatures** COSMIC mutational signature composition of HG008T graph-coordinate SNV predictions across WGS, PacBio, and ONT datasets. The stacked bars show the relative contribution of each inferred signature, and the adjacent bar plot shows the cosine similarity between each platform-specific signature profile and the HG008T benchmark profile. The relatively lower cosine similarity observed for HG008T may reflect the more heterogeneous and diffuse composition of the HG008T benchmark, which contains multiple substantial signature components. In contrast, COLO829T is dominated by a smaller set of signatures, particularly UV-associated signatures, resulting in stronger agreement between predicted and benchmark profiles. In addition, HG008T may depend more heavily on PoN-based filtering than COLO829T; therefore, the lack of PoN support for graph-node variants may lead to a greater loss of true somatic signal in HG008T.

#### Global .dat Header

| Field | Description | Type / Size | Value / Notes |
|---|---|---|---|
| magic | magic string | char[6] | b"MYFMT\x01" |
| version_major | major version | uint8_t | 0 |
| version_minor | minor version | uint8_t | 3 (cigar padded to node_length) |
| block_count | number of node blocks | uint32_t | count of blocks |
| reserved | reserved | char[16] | all 0x00 |
| | | | Total header size = 28 bytes |

#### Per-node Block Header

| Field | Description | Type / Size | Example |
|---|---|---|---|
| node_id | Graph node id | uint32_t | 123456 |
| n_records | Number of records in this block | uint32_t | 57 |
| flags | Reserved flags | uint16_t | 0 |
| node_length | Node sequence length | uint32_t | 143 |
| | | | Block header size = 14 bytes |

#### Segment Record Layout

| Field | Description | Type / Size | Notes |
|---|---|---|---|
| offset | alignment offset on node | int16_t (2B) | mapping.position.offset |
| seq | read bases | char[L] | padded/truncated to L |
| bq | base qualities | char[L] | Phred ASCII, padded |
| cigar | CIGAR string | char[L] | ASCII, padded/truncated |

| Field | Description | Type / Size | Notes |
| --- | --- | --- | --- |
| rq | mapping quality (MAPQ) | int16_t (2B) | alignment.mapping_quality |
| strand | strand | char (1B) | '+' or '-' |
| | | | Record size = 5 + 3L bytes |

#### .idx Index File Layout

| Field | Description | Type / Size | Notes |
|---|---|---|---|
| count | number of entries | uint32_t | file starts with this |
| node_id | node id | uint32_t | |
| offset | block offset in .dat | uint64_t | points to block header |
| block_size | block size in bytes | uint32_t | header + records |
| n_records | records in block | uint32_t | |
| flags | reserved | uint16_t | 0 |
| node_length | node length L | uint32_t | new index only (26B entry) |

**Supplementary Table 1. Node Pile Up (NPU) binary format specification.**

The table summarizes the binary layout of the NPU .dat and .idx files used to store graph-aligned read pileups. The .dat file contains a global header, per-node block headers and fixed-length segment records encoding node offsets, read bases, base qualities, CIGAR strings, mapping quality and strand information. The .idx file maps graph node IDs to the corresponding block offsets and sizes in the .dat file, enabling efficient random access to node-level pileup records.

##### Supplement Table 2, 3

| Dataset | Platform | Purity | Variant type | Caller | TP | FP | FN | Precision | Recall | F1 |
|---|---|---|---|---|---|---|---|---|---|---|
| **COLO829T** | WGS | P25 | SNV | Pansoma | 1478 | 242 | 965 | 0.8593 | 0.605 | 0.7101 |
| **COLO829T** | WGS | P50 | SNV | Pansoma | 1579 | 198 | 864 | 0.8886 | 0.6463 | 0.7483 |
| **COLO829T** | WGS | P75 | SNV | Pansoma | 1592 | 231 | 851 | 0.8733 | 0.6517 | 0.7464 |
| **COLO829T** | WGS | P100 | SNV | Pansoma | 1455 | 314 | 988 | 0.8225 | 0.5956 | 0.6909 |
| **COLO829T** | WGS | P25 | SNV | DeepSomatic-TO | 1799 | 8560 | 644 | 0.1737 | 0.7364 | 0.281 |
| **COLO829T** | WGS | P50 | SNV | DeepSomatic-TO | 1930 | 8387 | 513 | 0.1871 | 0.79 | 0.3025 |
| **COLO829T** | WGS | P75 | SNV | DeepSomatic-TO | 2019 | 8005 | 424 | 0.2014 | 0.8264 | 0.3239 |
| **COLO829T** | WGS | P100 | SNV | DeepSomatic-TO | 2067 | 5871 | 376 | 0.2604 | 0.8461 | 0.3982 |
| **COLO829T** | WGS | P25 | SNV | ClairS-TO | 1534 | 2129 | 909 | 0.4188 | 0.6279 | 0.5025 |
| **COLO829T** | WGS | P50 | SNV | ClairS-TO | 1697 | 1957 | 746 | 0.4644 | 0.6946 | 0.5567 |
| **COLO829T** | WGS | P75 | SNV | ClairS-TO | 1860 | 1814 | 583 | 0.5063 | 0.7614 | 0.6081 |
| **COLO829T** | WGS | P100 | SNV | ClairS-TO | 1890 | 2808 | 553 | 0.4023 | 0.7736 | 0.5293 |
| **COLO829T** | PacBio | P25 | SNV | Pansoma | 1340 | 417 | 1103 | 0.7627 | 0.5485 | 0.6381 |

| | | | | | | | | | | |
|---|---|---|---|---|---|---|---|---|---|---|
| **COLO829T** | PacBio | P50 | SNV | Pansoma | 1340 | 439 | 1103 | 0.7532 | 0.5485 | 0.6348 |
| **COLO829T** | PacBio | P75 | SNV | Pansoma | 1344 | 382 | 1099 | 0.7787 | 0.5501 | 0.6448 |
| **COLO829T** | PacBio | P100 | SNV | Pansoma | 1609 | 237 | 834 | 0.8716 | 0.6586 | 0.7503 |
| **COLO829T** | PacBio | P25 | SNV | DeepSomatic-TO | 1939 | 3474 | 504 | 0.3582 | 0.7937 | 0.4936 |
| **COLO829T** | PacBio | P50 | SNV | DeepSomatic-TO | 2069 | 3098 | 374 | 0.4004 | 0.8469 | 0.5438 |
| **COLO829T** | PacBio | P75 | SNV | DeepSomatic-TO | 2067 | 2642 | 376 | 0.4389 | 0.8461 | 0.578 |
| **COLO829T** | PacBio | P100 | SNV | DeepSomatic-TO | 1374 | 952 | 1069 | 0.5907 | 0.5624 | 0.5762 |
| **COLO829T** | PacBio | P25 | SNV | ClairS-TO | 1605 | 1713 | 838 | 0.4837 | 0.657 | 0.5572 |
| **COLO829T** | PacBio | P50 | SNV | ClairS-TO | 1768 | 1683 | 675 | 0.5123 | 0.7237 | 0.5999 |
| **COLO829T** | PacBio | P75 | SNV | ClairS-TO | 1799 | 1461 | 644 | 0.5518 | 0.7364 | 0.6309 |
| **COLO829T** | PacBio | P100 | SNV | ClairS-TO | 1839 | 772 | 604 | 0.7043 | 0.7528 | 0.7277 |
| **COLO829T** | ONT | P25 | SNV | Pansoma | 1424 | 669 | 1019 | 0.6804 | 0.5829 | 0.6279 |
| **COLO829T** | ONT | P50 | SNV | Pansoma | 1477 | 701 | 966 | 0.6781 | 0.6046 | 0.6393 |
| **COLO829T** | ONT | P75 | SNV | Pansoma | 1444 | 607 | 999 | 0.704 | 0.5911 | 0.6426 |
| **COLO829T** | ONT | P100 | SNV | Pansoma | 1405 | 313 | 1038 | 0.8178 | 0.5751 | 0.6753 |
| **COLO829T** | ONT | P25 | SNV | DeepSomatic-TO | 1682 | 3975 | 761 | 0.2973 | 0.6885 | 0.4153 |
| **COLO829T** | ONT | P50 | SNV | DeepSomatic-TO | 1854 | 4042 | 589 | 0.3145 | 0.7589 | 0.4447 |
| **COLO829T** | ONT | P75 | SNV | DeepSomatic-TO | 1873 | 3781 | 570 | 0.3313 | 0.7667 | 0.4626 |
| **COLO829T** | ONT | P100 | SNV | DeepSomatic-TO | 1721 | 2550 | 722 | 0.403 | 0.7045 | 0.5127 |
| **COLO829T** | ONT | P25 | SNV | ClairS-TO | 1412 | 1678 | 1031 | 0.457 | 0.578 | 0.5104 |
| **COLO829T** | ONT | P50 | SNV | ClairS-TO | 1639 | 1639 | 804 | 0.5 | 0.6709 | 0.573 |
| **COLO829T** | ONT | P75 | SNV | ClairS-TO | 1682 | 1379 | 761 | 0.5495 | 0.6885 | 0.6112 |
| **COLO829T** | ONT | P100 | SNV | ClairS-TO | 1691 | 828 | 752 | 0.6713 | 0.6922 | 0.6816 |
| **COLO829T** | WGS | P50 | INDEL | Pansoma | 22 | 167 | 109 | 0.1164 | 0.1679 | 0.1375 |
| **COLO829T** | WGS | P75 | INDEL | Pansoma | 28 | 122 | 103 | 0.1867 | 0.2137 | 0.1993 |
| **COLO829T** | WGS | P100 | INDEL | Pansoma | 29 | 104 | 102 | 0.218 | 0.2214 | 0.2197 |
| **COLO829T** | WGS | P25 | INDEL | DeepSomatic-TO | 24 | 3343 | 107 | 0.0071 | 0.1832 | 0.0137 |
| **COLO829T** | WGS | P50 | INDEL | DeepSomatic-TO | 35 | 2990 | 96 | 0.0116 | 0.2672 | 0.0222 |
| **COLO829T** | WGS | P75 | INDEL | DeepSomatic-TO | 34 | 2025 | 97 | 0.0165 | 0.2595 | 0.0311 |
| **COLO829T** | WGS | P100 | INDEL | DeepSomatic-TO | 28 | 1010 | 103 | 0.027 | 0.2137 | 0.0479 |
| **COLO829T** | WGS | P25 | INDEL | ClairS-TO | 6 | 142 | 125 | 0.0405 | 0.0458 | 0.043 |
| **COLO829T** | WGS | P50 | INDEL | ClairS-TO | 14 | 136 | 117 | 0.0933 | 0.1069 | 0.0996 |
| **COLO829T** | WGS | P75 | INDEL | ClairS-TO | 13 | 99 | 118 | 0.1161 | 0.0992 | 0.107 |
| **COLO829T** | WGS | P100 | INDEL | ClairS-TO | 19 | 194 | 112 | 0.0892 | 0.145 | 0.1105 |

| | | | | | | | | | | |
|---|---|---|---|---|---|---|---|---|---|---|
| **COLO829T** | PacBio | P50 | INDEL | Pansoma | 8 | 54 | 123 | 0.1290 | 0.0611 | 0.0829 |
| **COLO829T** | PacBio | P75 | INDEL | Pansoma | 13 | 44 | 118 | 0.2281 | 0.0992 | 0.1383 |
| **COLO829T** | PacBio | P100 | INDEL | Pansoma | 18 | 64 | 113 | 0.2195 | 0.1374 | 0.1690 |
| **COLO829T** | PacBio | P25 | INDEL | DeepSomatic-TO | 9 | 223 | 122 | 0.0388 | 0.0687 | 0.0496 |
| **COLO829T** | PacBio | P50 | INDEL | DeepSomatic-TO | 17 | 188 | 114 | 0.0829 | 0.1298 | 0.1012 |
| **COLO829T** | PacBio | P75 | INDEL | DeepSomatic-TO | 13 | 113 | 118 | 0.1032 | 0.0992 | 0.1012 |
| **COLO829T** | PacBio | P100 | INDEL | DeepSomatic-TO | 12 | 39 | 119 | 0.2353 | 0.0916 | 0.1319 |
| **COLO829T** | PacBio | P25 | INDEL | ClairS-TO | 3 | 108 | 128 | 0.027 | 0.0229 | 0.0248 |
| **COLO829T** | PacBio | P50 | INDEL | ClairS-TO | 11 | 79 | 120 | 0.1222 | 0.084 | 0.0995 |
| **COLO829T** | PacBio | P75 | INDEL | ClairS-TO | 14 | 48 | 117 | 0.2258 | 0.1069 | 0.1451 |
| **COLO829T** | PacBio | P100 | INDEL | ClairS-TO | 14 | 46 | 117 | 0.2333 | 0.1069 | 0.1466 |
| **COLO829T** | ONT | P50 | INDEL | Pansoma | 5 | 886 | 126 | 0.0056 | 0.0382 | 0.0098 |
| **COLO829T** | ONT | P75 | INDEL | Pansoma | 7 | 708 | 124 | 0.0098 | 0.0534 | 0.0165 |
| **COLO829T** | ONT | P100 | INDEL | Pansoma | 8 | 517 | 123 | 0.0152 | 0.0611 | 0.0244 |
| **COLO829T** | ONT | P25 | INDEL | DeepSomatic-TO | 3 | 178 | 128 | 0.0166 | 0.0229 | 0.0192 |
| **COLO829T** | ONT | P50 | INDEL | DeepSomatic-TO | 6 | 165 | 125 | 0.0351 | 0.0458 | 0.0397 |
| **COLO829T** | ONT | P75 | INDEL | DeepSomatic-TO | 3 | 126 | 128 | 0.0233 | 0.0229 | 0.0231 |
| **COLO829T** | ONT | P100 | INDEL | DeepSomatic-TO | 2 | 80 | 129 | 0.0244 | 0.0153 | 0.0188 |
| **COLO829T** | ONT | P25 | INDEL | ClairS-TO | 1 | 97 | 130 | 0.0102 | 0.0076 | 0.0087 |
| **COLO829T** | ONT | P50 | INDEL | ClairS-TO | 8 | 88 | 123 | 0.0833 | 0.0611 | 0.0705 |
| **COLO829T** | ONT | P75 | INDEL | ClairS-TO | 13 | 73 | 118 | 0.1512 | 0.0992 | 0.1198 |
| **COLO829T** | ONT | P100 | INDEL | ClairS-TO | 12 | 108 | 119 | 0.1 | 0.0916 | 0.0956 |
| **HG008T** | WGS | Tumor-only | SNV | Pansoma | 565 | 549 | 137 | 0.5072 | 0.8048 | 0.6222 |
| **HG008T** | WGS | Tumor-only | SNV | ClairS-TO | 603 | 1676 | 99 | 0.2646 | 0.859 | 0.4046 |
| **HG008T** | WGS | Tumor-only | SNV | DeepSomatic-TO | 639 | 4838 | 63 | 0.1167 | 0.9103 | 0.2068 |
| **HG008T** | PacBio | Tumor-only | SNV | Pansoma | 533 | 486 | 169 | 0.5231 | 0.7593 | 0.6194 |
| **HG008T** | PacBio | Tumor-only | SNV | ClairS-TO | 601 | 1131 | 101 | 0.347 | 0.8561 | 0.4938 |
| **HG008T** | PacBio | Tumor-only | SNV | DeepSomatic-TO | 572 | 1758 | 130 | 0.2455 | 0.8148 | 0.3773 |
| **HG008T** | ONT | Tumor-only | SNV | Pansoma | 581 | 568 | 121 | 0.5057 | 0.8276 | 0.6278 |
| **HG008T** | ONT | Tumor-only | SNV | ClairS-TO | 593 | 1302 | 109 | 0.3129 | 0.8447 | 0.4567 |
| **HG008T** | ONT | Tumor-only | SNV | DeepSomatic-TO | 618 | 7549 | 84 | 0.0757 | 0.8803 | 0.1394 |

| | | | | | | | | | | |
|---|---|---|---|---|---|---|---|---|---|---|
| **HG008T** | WGS | Tumor-only | INDEL | Pansoma | 122 | 639 | 513 | 0.1603 | 0.1921 | 0.1748 |
| **HG008T** | WGS | Tumor-only | INDEL | ClairS-TO | 50 | 128 | 585 | 0.2809 | 0.0787 | 0.123 |
| **HG008T** | WGS | Tumor-only | INDEL | DeepSomatic-TO | 108 | 2450 | 527 | 0.0422 | 0.1701 | 0.0676 |
| **HG008T** | PacBio | Tumor-only | INDEL | Pansoma | 82 | 457 | 553 | 0.1521 | 0.1291 | 0.1397 |
| **HG008T** | PacBio | Tumor-only | INDEL | ClairS-TO | 47 | 83 | 588 | 0.3615 | 0.074 | 0.1229 |
| **HG008T** | PacBio | Tumor-only | INDEL | DeepSomatic-TO | 23 | 150 | 612 | 0.1329 | 0.0362 | 0.0569 |
| **HG008T** | ONT | Tumor-only | INDEL | Pansoma | 116 | 1684 | 519 | 0.0644 | 0.1827 | 0.0953 |
| **HG008T** | ONT | Tumor-only | INDEL | ClairS-TO | 47 | 92 | 588 | 0.3381 | 0.074 | 0.1214 |
| **HG008T** | ONT | Tumor-only | INDEL | DeepSomatic-TO | 35 | 3210 | 600 | 0.0108 | 0.0551 | 0.018 |

| **Dataset** | **Platform** | **Benchmark** | **Variant type** | **Caller** | **VAF bin** | **TP (precision evaluation)** | **FP (precision evaluation)** | **TP (recall evaluation)** | **FN (recall evaluation)** | **Precision** | **Recall** | **F1** |
|---|---|---|---|---|---|---|---|---|---|---|---|---|
| **HapMap** | WGS | Low-VAF mixture | SNV | Pansoma | 1–16.5% | 174082 | 29645 | 90405 | 5988 | 0.8545 | 0.9379 | 0.8942 |
| **HapMap** | WGS | Low-VAF mixture | SNV | Pansoma | 1–2% | 31562 | 7033 | 15035 | 2112 | 0.8178 | 0.8768 | 0.8463 |
| **HapMap** | WGS | Low-VAF mixture | SNV | Pansoma | 2–5% | 73372 | 14050 | 52610 | 1852 | 0.8393 | 0.9660 | 0.8982 |
| **HapMap** | WGS | Low-VAF mixture | SNV | Pansoma | 5–10% | 59989 | 6072 | 21178 | 1463 | 0.9081 | 0.9354 | 0.9215 |
| **HapMap** | WGS | Low-VAF mixture | SNV | Pansoma | 10–16.5% | 9159 | 2490 | 1582 | 561 | 0.7862 | 0.7382 | 0.7614 |
| **HapMap** | WGS | Low-VAF mixture | SNV | Mutect2 | 1–16.5% | 220686 | 10135 | 65618 | 30775 | 0.9561 | 0.6807 | 0.7953 |
| **HapMap** | WGS | Low-VAF mixture | SNV | Mutect2 | 1–2% | 55716 | 2523 | 10561 | 6586 | 0.9567 | 0.6159 | 0.7494 |
| **HapMap** | WGS | Low-VAF mixture | SNV | Mutect2 | 2–5% | 54959 | 2895 | 39077 | 15385 | 0.9500 | 0.7175 | 0.8175 |
| **HapMap** | WGS | Low-VAF mixture | SNV | Mutect2 | 5–10% | 83805 | 3221 | 14939 | 7702 | 0.9630 | 0.6598 | 0.7831 |
| **HapMap** | WGS | Low-VAF mixture | SNV | Mutect2 | 10–16.5% | 26206 | 1496 | 1041 | 1102 | 0.9460 | 0.4858 | 0.6419 |
| **HapMap** | WGS | Low-VAF mixture | INDEL | Pansoma | 1–16.5% | 33131 | 15496 | 22697 | 53848 | 0.6813 | 0.2965 | 0.4132 |
| **HapMap** | WGS | Low-VAF mixture | INDEL | Pansoma | 1–2% | 12218 | 6404 | 3760 | 13093 | 0.6561 | 0.2231 | 0.3330 |

| | | | | | | | | | | | |
|---|---|---|---|---|---|---|---|---|---|---|---|
| **HapMap** | WGS | Low-VAF mixture | INDEL | Pansoma | 2–5% | 12736 | 5232 | 10002 | 17624 | 0.7088 | 0.3621 | 0.4793 |
| **HapMap** | WGS | Low-VAF mixture | INDEL | Pansoma | 5–10% | 7000 | 2378 | 8278 | 17102 | 0.7464 | 0.3262 | 0.4540 |
| **HapMap** | WGS | Low-VAF mixture | INDEL | Pansoma | 10–16.5% | 1177 | 1482 | 657 | 6029 | 0.4426 | 0.0983 | 0.1609 |
| **HapMap** | WGS | Low-VAF mixture | INDEL | Mutect2 | 1–16.5% | 14586 | 2481 | 11218 | 65438 | 0.8546 | 0.1463 | 0.2498 |
| **HapMap** | WGS | Low-VAF mixture | INDEL | Mutect2 | 1–2% | 3181 | 443 | 1793 | 15087 | 0.8778 | 0.1062 | 0.1895 |
| **HapMap** | WGS | Low-VAF mixture | INDEL | Mutect2 | 2–5% | 3651 | 749 | 4411 | 23259 | 0.8298 | 0.1594 | 0.2674 |
| **HapMap** | WGS | Low-VAF mixture | INDEL | Mutect2 | 5–10% | 5855 | 880 | 3768 | 21646 | 0.8693 | 0.1483 | 0.2534 |
| **HapMap** | WGS | Low-VAF mixture | INDEL | Mutect2 | 10–16.5% | 1899 | 409 | 1246 | 5446 | 0.8228 | 0.1862 | 0.3037 |

**Supplementary Tables 2 and 3. Benchmarking performance across COLO829T, HG008T and HapMap datasets.**

Supplementary Table 2 summarizes SNV and INDEL calling performance for Pansoma, ClairS-TO and DeepSomatic-TO across COLO829T and HG008T datasets, including WGS, PacBio HiFi and ONT platforms, COLO829T tumor-purity series and HG008T tumor-only evaluations. Supplementary Table 3 summarizes low-VAF HapMap WGS benchmark performance for Pansoma and Mutect2 across SNV and INDEL VAF bins from 1–2% to 10–16.5%, together with the overall 1–16.5% range. For each benchmark, true positives, false positives, false negatives, precision, recall and F1 score are reported; in the HapMap table, TP counts are separated for precision and recall evaluations because call-set and truth-set VAF stratifications were evaluated independently.

| category | metric | WGS | ONT | PacBio |
|---|---|---|---|---|
| **tool_counts** | Pansoma-TO | 1114 | 1149 | 1019 |
| **tool_counts** | ClairS-TO | 2279 | 1895 | 1732 |
| **tool_counts** | DeepSomatic-TO | 5477 | 8169 | 2331 |
| **tool_counts** | Ground Truth | 702 | 702 | 702 |
| **venn_regions** | Pansoma-TO only | 65 | 76 | 52 |
| **venn_regions** | ClairS-TO only | 446 | 76 | 146 |
| **venn_regions** | DeepSomatic-TO only | 3595 | 6297 | 800 |
| **venn_regions** | Ground Truth only | 58 | 59 | 65 |
| **venn_regions** | Pansoma-TO&ClairS-TO only | 12 | 14 | 47 |
| **venn_regions** | Pansoma-TO&DeepSomatic-TO only | 25 | 42 | 21 |
| **venn_regions** | Pansoma-TO&Ground Truth only | 0 | 9 | 8 |

| | | | | |
|---|---|---|---|---|
| **venn_regions** | ClairS-TO&DeepSomatic-TO only | 771 | 776 | 572 |
| **venn_regions** | ClairS-TO&Ground Truth only | 3 | 4 | 20 |
| **venn_regions** | DeepSomatic-TO&Ground Truth only | 21 | 12 | 10 |
| **venn_regions** | Pansoma-TO&ClairS-TO&DeepSomatic-TO only | 447 | 436 | 366 |
| **venn_regions** | Pansoma-TO&ClairS-TO&Ground Truth only | 2 | 12 | 37 |
| **venn_regions** | Pansoma-TO&DeepSomatic-TO&Ground Truth only | 20 | 29 | 18 |
| **venn_regions** | ClairS-TO&DeepSomatic-TO&Ground Truth only | 55 | 46 | 74 |
| **venn_regions** | Pansoma-TO&ClairS-TO&DeepSomatic-TO&Ground Truth | 543 | 531 | 470 |
| **key_ratios** | DeepSomatic-TO covers Pansoma-TO | 92.9% | 90.3% | 85.9% |
| **key_ratios** | ClairS-TO covers Pansoma-TO | 90.1% | 86.4% | 90.3% |
| **key_ratios** | Pansoma-TO captures Ground Truth | 80.5% | 82.8% | 75.9% |
| **key_ratios** | ClairS-TO captures Ground Truth | 85.9% | 84.5% | 85.6% |
| **key_ratios** | DeepSomatic-TO captures Ground Truth | 91.0% | 88.0% | 81.5% |
| **key_ratios** | All callers share Ground Truth | 77.4% | 75.6% | 67.0% |

| category | metric | WGS | ONT | PacBio |
|---|---|---|---|---|
| **tool_counts** | Pansoma-TO | 1769 | 1718 | 1757 |
| **tool_counts** | ClairS-TO | 4698 | 2519 | 2611 |
| **tool_counts** | DeepSomatic-TO | 7938 | 4273 | 2326 |
| **tool_counts** | Ground Truth | 2443 | 2443 | 2443 |
| **venn_regions** | Pansoma-TO only | 129 | 116 | 299 |
| **venn_regions** | ClairS-TO only | 1885 | 193 | 487 |
| **venn_regions** | DeepSomatic-TO only | 4944 | 1910 | 730 |
| **venn_regions** | Ground Truth only | 337 | 455 | 266 |
| **venn_regions** | Pansoma-TO&ClairS-TO only | 7 | 23 | 72 |
| **venn_regions** | Pansoma-TO&DeepSomatic-TO only | 11 | 30 | 9 |
| **venn_regions** | Pansoma-TO&Ground Truth only | 3 | 60 | 47 |
| **venn_regions** | ClairS-TO&DeepSomatic-TO only | 749 | 468 | 176 |
| **venn_regions** | ClairS-TO&Ground Truth only | 21 | 73 | 188 |
| **venn_regions** | DeepSomatic-TO&Ground Truth only | 161 | 151 | 263 |
| **venn_regions** | Pansoma-TO&ClairS-TO&DeepSomatic-TO only | 167 | 144 | 37 |
| **venn_regions** | Pansoma-TO&ClairS-TO&Ground Truth only | 15 | 134 | 568 |
| **venn_regions** | Pansoma-TO&DeepSomatic-TO&Ground Truth only | 52 | 86 | 28 |
| **venn_regions** | ClairS-TO&DeepSomatic-TO&Ground Truth only | 469 | 359 | 386 |
| **venn_regions** | Pansoma-TO&ClairS-TO&DeepSomatic-TO&Ground Truth | 1385 | 1125 | 697 |
| **key_ratios** | DeepSomatic-TO covers Pansoma-TO | 91.3% | 80.6% | 43.9% |
| **key_ratios** | ClairS-TO covers Pansoma-TO | 89.0% | 83.0% | 78.2% |
| **key_ratios** | Pansoma-TO captures Ground Truth | 59.6% | 57.5% | 54.9% |
| **key_ratios** | ClairS-TO captures Ground Truth | 77.4% | 69.2% | 75.3% |

| | | | | |
|---|---|---|---|---|
| **key_ratios** | DeepSomatic-TO captures Ground Truth | 84.6% | 70.4% | 56.2% |
| **key_ratios** | All callers share Ground Truth | 56.7% | 46.0% | 28.5% |

**Supplementary Table 4 and 5. Callers overlap counts for PASS-only SNV calls.**

The table 4a and 4b are HG008T and COLO829T respectively. For each sample, the table reports total PASS SNV calls from Pansoma-TO, ClairS-TO, DeepSomatic-TO, and the Ground Truth set, followed by mutually exclusive overlap regions among the four sets across WGS, ONT, and PacBio data. Key ratios summarize the fraction of Pansoma-TO calls covered by ClairS-TO or DeepSomatic-TO, and the fraction of Ground Truth variants captured by each caller or shared by all callers.

| Dataset | Platform | Group | Linear count | Graph-coordinate count | Total count | Graph-coordinate fraction (%) |
|---|---|---|---|---|---|---|
| **SMaHT** | WGS | P25 | 1720 | 52 | 1772 | 2.93 |
| **SMaHT** | WGS | P50 | 1777 | 53 | 1830 | 2.90 |
| **SMaHT** | WGS | P75 | 1823 | 60 | 1883 | 3.19 |
| **SMaHT** | WGS | P100 | 1769 | 57 | 1826 | 3.12 |
| **SMaHT** | PacBio | P25 | 1757 | 96 | 1853 | 5.18 |
| **SMaHT** | PacBio | P50 | 1779 | 94 | 1873 | 5.02 |
| **SMaHT** | PacBio | P75 | 1726 | 86 | 1812 | 4.75 |
| **SMaHT** | PacBio | P100 | 1846 | 94 | 1940 | 4.85 |
| **SMaHT** | ONT | P25 | 2093 | 403 | 2496 | 16.15 |
| **SMaHT** | ONT | P50 | 2178 | 287 | 2465 | 11.64 |
| **SMaHT** | ONT | P75 | 2051 | 328 | 2379 | 13.79 |
| **SMaHT** | ONT | P100 | 1718 | 530 | 2248 | 23.58 |
| **HG008** | WGS | No HG008N | 1114 | 57 | 1171 | 4.87 |
| **HG008** | WGS | HG008N added | 641 | 44 | 685 | 6.42 |

**Supplementary Table 6. Chromosome 1 variant counts stratified by linear and graph-coordinate representation.**

Summary of chr1 variant counts detected on the GRCh38 linear reference and graph-coordinate paths across SMaHT WGS, PacBio HiFi and ONT datasets, as well as HG008 WGS analyses with and without HG008N added. Total counts were calculated as the sum of linear and graph-coordinate variants, and graph-coordinate fractions indicate the proportion of variants represented outside the linear-reference count.
